# High-throughput RNA sequencing from paired lesional- and non-lesional skin reveals major alterations in the psoriasis circRNAome

**DOI:** 10.1101/581066

**Authors:** Liviu-Ionut Moldovan, Thomas Birkballe Hansen, Morten Trillingsgaard Venø, Trine Line Hauge Okholm, Thomas Levin Andersen, Henrik Hager, Lars Iversen, Jørgen Kjems, Claus Johansen, Lasse Sommer Kristensen

## Abstract

**Background:** Psoriasis is a chronic inflammatory skin disease characterized by hyperproliferation and abnormal differentiation of keratinocytes. It is one of the most prevalent chronic inflammatory skin condition in adults worldwide, with a considerable negative impact on quality of life. Circular RNAs (circRNAs) are a recently identified type of non-coding RNA with diverse cellular functions related to their exceptional stability. In particular, some circRNAs can bind and regulate microRNAs (miRNAs), a group of RNAs that play a role in the pathogenesis of psoriasis. The aim of this study was to characterize the circRNAome in psoriasis and to assess potential correlations to miRNA expression patterns.

**Results:** Using high-throughput RNA-sequencing (RNA-seq) and NanoString nCounter technology, we found a substantial down-regulation of circRNA expression in lesional skin compared to non-lesional skin from psoriasis patients. We saw that this mainly applies to the epidermis by analyzing laser capture microdissected tissues and by RNA chromogenic *in situ* hybridization (CISH). We also found that the majority of the circRNAs were downregulated independent of their corresponding linear host genes. The observed downregulation of circRNAs in psoriasis was not due to altered expression levels of factors known to affect circRNA biogenesis, nor because lesional skin contained an increased number of inflammatory cells such as lymphocytes. Finally, we saw that the overall differences in available miRNA binding sites on the circRNAs between lesional and non-lesional skin did not correlate with differences in miRNA expression patterns.

**Conclusions:** We have performed the first genome-wide circRNA profiling of paired lesional and non-lesional skin from psoriasis patients and revealed that circRNAs are much less abundant in the lesional samples. Whether this is a cause or a consequence of the disease remains to be revealed, however, we found no evidence that the loss of miRNA binding sites on the circRNAs could explain differences in miRNA expression reported between lesional and non-lesional skin.

## Background

Psoriasis is one of the most common chronic inflammatory skin conditions, with 1-3% of the adult population affected worldwide [1]. It is defined by a pronounced hyperproliferation and deficient terminal differentiation of the keratinocytes. Moreover, a complex interplay between different cell types (e.g. T cells and dendritic cells) and a variety of cytokines are known to contribute in the development of psoriasis. The pathogenesis is also based on a complex synergy between genetic predisposition, major histocompatibility alleles, and a variety of environmental triggers [2]. From the molecular point of view, however, the mechanisms responsible for the interactions between keratinocytes and inflammatory cells infiltrating the epidermis are still not fully understood. The analysis of the molecular backdrop of psoriasis has described numerous disease-associated genes and proteins with abnormal expression patterns [3], but little is known about the regulatory pathways responsible for this aberrant expression. Recent findings suggest that non-coding RNAs such as microRNAs (miRNAs) and long non-coding RNAs (lncRNAs) could be involved in the pathogenesis of psoriasis by influencing protein expression and function in both keratinocytes and inflammatory cells [4-8].

Recent discoveries have let to the identification of a subclass of non-coding RNA, the circular RNAs (circRNAs), which are formed by a backsplicing event linking the 3′ end of an exon to the 5′ end of the same or an upstream exon [9-11]. Most circRNAs are expressed from known protein coding genes and in human cells more unique circRNAs than genes have been annotated [12]. They typically reside in the cytoplasm, are highly stable, evolutionary conserved, and often abnormally expressed in a variety of human diseases, most notably in cancer [13, 14]. Backsplicing is facilitated by the presence of homologous inverted repeats (e.g. inverted Alu elements (IAE)) in the regions flanking the involved exons [9, 14-17]. These inverted elements can base-pair to create a loop structure of the nascent RNA, bringing the splice sites involved in backsplicing into close proximity. In support of this biogenesis mechanism, the RNA helicase DHX9 and the RNA editing enzyme ADAR1 were shown to bind double stranded RNA formed by IAEs and suppress circular RNA production [15-17]. Conversely, the splicing factors QKI, FUS, and HNRNPL endorse the production of circRNA by binding to flanking motifs and forming dimers, which bring the splice sites involved in backsplicing into close proximity [18-20].

The functions of the majority of the annotated circRNAs remain to be disclosed; however, evidence suggests that at least some function as miRNA sponges [10, 21-23]. miRNAs are a class of 21-23 nt RNAs involved in gene-regulation at the post-transcriptional level, mainly by directly interacting with the 3’ untranslated region of mRNAs. This results in repression of translation by mRNA destabilization or degradation [24]. The circular RNA sponge for miR-7 (ciRS-7), also known as CDR1as, is the most studied circular RNA sponge. It contains more than 70 conserved binding sites for miR-7 [21] but it is still debated whether it acts as an inhibitor, a protector or even as a transporter of miR-7 [25, 26]. However, ciRS-7 is not a typical circRNA and the majority do not harbor more miRNA binding sites than expected by chance [27]. Therefore, it is still widely debated whether the ability to acts as a miRNA sponge is a general feature of circRNAs.

The genome-wide landscape of circRNA expression (the circRNAome) has not previously been revealed in paired primary skin samples from psoriasis patients. We profiled the circRNAome in paired lesional- and non-lesional skin using high-throughput RNA-sequencing (RNA-seq) and investigated possible mechanisms behind the major alterations observed. We further investigated circRNA expression patterns in individual cell types within the tissues and explored how the dramatic changes observed in the circRNAome impact genome-wide miRNA expression patterns.

## Materials and Methods

### Patient samples

Two cohorts of patients of Caucasian origin diagnosed with psoriasis vulgaris were recruited for this study. The first cohort consisted of six subjects (average age 42.8 years, range 35–52 years, 1 woman and 5 men). The mean baseline PASI was 16.2 (12-20.8). PASI for patient 6 was not available. The second cohort consisted of 14 subjects (average age 44.9 years, range 26-62 years, 5 women and 9 men). The mean baseline PASI was 25.6 (17.2-51.8). None of the participants had used any systemically immunosuppressive medications for four weeks and none had received local treatment at the site of biopsies for two weeks before study participation. Four-millimeter punch biopsies were obtained from lesional and non-lesional psoriatic skin taken from the center of a plaque from patients with moderate to severe chronic stable plaque psoriasis from either the upper or the lower extremities. For each patient, biopsies were taken from only one anatomical site. Biopsies were taken as paired samples and the ones from non-lesional psoriatic skin were taken from the same body region as the ones from lesional psoriatic skin at a distance of at least five centimeters from a plaque. The biopsies were immediately snap-frozen in liquid nitrogen and stored until further use.

### Ethical approval

This study was conducted according to the Declaration of Helsinki Principles. Informed consent was obtained from each patient and permission from the national ethics committee was granted (1807975).

### RNA extraction

Punch biopsies from psoriatic patients were transferred to 1 ml of -80°C cold RNAlater-ICE (Ambion inc., Austin, TX). Samples were kept at -80°C until 24 h before RNA purification at which time they were transferred to -20°C. Upon RNA purification, biopsies were removed from

RNAlater-ICE and transferred to 175 μl of SV RNA Lysis Buffer added β-mercaptoethanol (SV Total RNA Isolation System; Promega, Madison, WI) and homogenized. RNA purification, including DNase treatment of the samples was completed according to manufacturer’s instructions (SV Total RNA Isolation System; Promega, Madison, WI). The RNA was stored at -80°C until further use.

### RNA-seq

One microgram of total RNA was rRNA depleted using the Ribo-Zero rRNA Removal Kit (Human, Mouse, Rat) (Epicentre, Madison, WI, USA) followed by a purification step using AMPure XP Beads (Beckman Coulter, Brea, CA, USA). Sequencing libraries were generated using the ScriptSeq v2 RNA-Seq Library Preparation Kit (Epicentre) using 12 PCR cycles for amplification. Purification was performed using AMPure XP Beads (Beckman Coulter). The final libraries were size selected (150-500 bp) on a Pippin Prep (Sage Science, Inc. Beverly, MA, USA) and quality controlled on the 2100 Bioanalyzer (Agilent Technologies, Santa Clara, CA, USA) and quantified using the KAPA library quantification kit (Kapa Biosystems, Wilmington, MA, USA). RNA-seq was performed on the HiSeq 4000 system (Illumina, San Diego, CA, USA) at the Beijing Genomics Institute (BGI) in Copenhagen using the 100 paired-end sequencing protocol with twelve samples pooled on one lane.

### circRNA quantification in RNA-seq data

Sequencing data from lesional- and non-lesional skin biopsies were quality filtered (Phred score 20) and adapter trimmed using Trim Galore. Filtered data were mapped to the human genome (hg19) using Bowtie2, mapping only unspliced reads. Unmapped reads were analyzed using a stringent version of the find_circ bioinformatics algorithm [28]. Filtered reads were also mapped to hg19 using Tophat2 and analyzed using CIRCexplorer [29]. All circRNA data analyses were based on the stringent version of the find_circ pipeline and circRNA candidates supported by at least five BSJ-spanning reads on average per sample were defined as high abundance circRNAs. Among these, all circRNA candidates not detected by CIRCexplorer were manually inspected to exclude obvious mapping artifacts as previously described [22]. Reads per million (RPM) refers to sequencing reads aligning across the particular BSJ divided by the total number of reads multiplied by one million. The circular-to-linear (CTL) ratios were defined as the number of reads spanning the particular BSJs divided by the average linear reads spanning the splice donor- and splice acceptor sites of the same BSJ plus one pseudo count (to avoid division by zero).

### mRNA quantification in RNA-seq data

Sequencing reads were quality-filtered, and adaptor-trimmed as described above. Filtered reads were mapped to hg19 using Tophat2 and featureCounts [30] was used to quantify the number of reads mapping to annotated genes from Ensembl gene definitions release 71. Differential expression analysis was performed using the DESeq2 R package.

### Analyses of IAEs

We used the UCSC Browser RepeatMasker track to extract the distance to nearest IAE flanking the BSJs of the circRNAs analyzed as previously described (https://www.biorxiv.org/content/early/2018/05/25/328104) and by only considering IAEs within the same subfamily (e.g. AluJ or AluS) as previously described [31].

### circRNA expression analyses using NanoString nCounter technology

A custom CodeSet of capture- and reporter probes was designed to target regions of 100 bp overlaying the BSJs of the top 50 most abundant circRNAs in the entire dataset (Supplementary Table 1). In addition, seven linear reference genes were included, which have previously been shown to be stable in lesional skin and non-lesional skin from patients with psoriasis [32]. Approximately 150 ng of total RNA from each sample was subjected to nCounter™ *SPRINT* (NanoString Technologies, Seattle, WA, USA) analysis according to the manufacturer’s instructions. The raw data were processed using the nSOLVER 3.0 software (NanoString Technologies); first, a background subtraction was performed using the max of negative controls, and then positive control normalization was performed using the geometric mean of all positive controls. Finally, a second normalization using the geometric mean of the three most stable linear reference genes (*ACTB, GUSB* and *RPL19*) was performed, before exporting the data to Excel (Microsoft Corporation, Redmond, WA, USA).

### Gene expression analyses of factors involved in circRNA biogenesis

Within the custom CodeSet for circRNA expression analysis, we also included capture- and reporter probes designed to target factors that are involved in circRNA biogenesis. These included *ADAR ADAR* [16], *DHX9* [17], *FUS* [20], *QKI* [19] and *HNRNPL* [18] (Supplementary Table 1). Data processing, including background subtraction and normalization, was performed as described above.

### miRNA expression analyses

The nCounter Human v3 miRNA panel (NanoString Technologies), which target 799 miRNAs, was used for miRNA profiling. One-hundred ng of total RNA from each sample was subjected to nCounter™ *SPRINT* (NanoString Technologies) analysis according to the manufacturer’s instructions. The raw data were processed using the nSOLVER 3.0 software (NanoString Technologies); first, a background subtraction was performed using the max of negative controls, and then positive control normalization was performed using the geometric mean of all positive controls. A second normalization was performed using the geometric mean of the best combination of any two miRs (hsa-let-7d-5p and hsa-miR-423-5p) as determined by the NormFinder algorithm [33], before exporting the data to Excel (Microsoft Corporation).

### Integrated analysis of miRNA- and circRNA data

To predict miRNA binding sites within the high abundance circRNAs, we first extracted the expected sequence of the mature circRNAs assuming splicing pattern similar to the host genes by which their share splice sites. Then, for all miRNAs analyzed, we determined the number of 8mer target sites present in the circRNA sequences. For each circRNA that was identified as having one or more miRNA binding sites, we calculated the ΔRPM values (mean circRNA RPM (lesional) -mean circRNA RPM (non-lesional)). Then, we performed a linear regression analysis of the sum of the ΔRPM values corresponding to each individual miRNA and either the fold change of each miRNA (mean miRNA counts (lesional)/mean miR counts (non-lesional)) or the absolute difference in expression level (mean miR counts (lesional) -mean miR counts (non-lesional)).

### Microdissections

After cutting the FFPE punch biopsies into 4 µm thick slices, they were mounted on membrane slides (Leica, Germany). The biopsies were deparaffinized for 30 s in Xylene, rehydrated in graded ethanol, stained for 2 s in hemalun (Mayers Hemalun, Merck, Germany) and washed in sterile water. After drying, 10 consecutive sections, epidermis and dermis were microdissected and collected in different tubes using a microdissection system (LMD 630, Leica Germany). RNA purification was done using the miRNeasy FFPE kit (Qiagen, Hilden, Germany).

### RNA chromogenic in situ hybridization (CISH)

The abundance of the circRNA, ciRS-7, in 10 lesional and 10 non-lesional skin biopsies was investigated by CISH using a modified version of the RNAScope 2.5 high-definition procedure (310035, Advanced Cell Diagnostics [ACD], Hayward, CA, USA), as previously described [34]. 3.5-µm-thick paraffin sections were pretreated and hybridized with 12 ZZ-pairs (Probe-Hs-CDR1-AS-No-XMm, 510711, ACD) targeting ciRS-7 overnight. The ZZ-pairs binding ciRS-7 were detected using seven amplification steps, including a Tyramid Signal Amplification step (TSA-DIG; NEL748001KT, PerkinElmer, Skovlunde, Denmark) labeled with alkaline phosphatase–conjugated sheep anti-DIG FAB fragments (11093274910, Roche, Basel, Switzerland), before visualized with Liquid Permanent Red (DAKO, Glostrup, Denmark) and counterstained with Mayer’s hematoxylin.

### Statistical analyses

All statistical tests concerning the circRNA analyses were performed using Prism 7 (GraphPad, La Jolla, CA, USA). Comparisons between the expression levels of the high abundance circRNAs and miRNAs in the lesional- and normal skin were done using Wilcoxon matched-pairs signed rank tests, as the data were not normally distributed according to the D’Agostino & Pearson normality test. Volcano plots were generated by one unpaired t test per circRNA individually without assuming consistent standard deviation and without correction for multiple testing. Linear regression was used to assess the potential correlation between fold changes in RPM and fold changes in CTL ratios employing an F test to investigate if the slope was significantly non-zero. Assessment of differences in expression levels of *ADAR, QKI, FUS, DHX9* and *HNRNPL* as well as T-cell-specific genes between the lesional- and non-lesional skin was done using paired t tests. Assessing the distance range to inverted Alu elements between circRNAs downregulated > two fold in lesional skin and the remaining circRNAs was done using Chi-squared tests and Mann Whitney tests. Analyses of circRNA and mRNA expression in the microdissected samples were done using unpaired t tests with Welch’s corrections, as data were not available for all sample pairs. All *P*-values were two-tailed and considered significant if□<□0.05.

## Results

### Identification and characterization of circular RNAs in lesional- and non-lesional skin biopsies

In order to perform an unbiased genome-wide profiling of circRNAs in psoriasis, we performed RNA-seq of ribosomal RNA-depleted total RNA from paired lesional- (n=6) and non-lesional (n=6) skin biopsies. We detected 2,066 and 2,626 unique circRNAs supported by at least two BSJ-spanning reads in a single sample in the lesional- and non-lesional skin biopsies, respectively, using a stringent version of the find_circ bioinformatics algorithm [28]. However, in order to ensure that we are looking at bona fide circRNAs with reasonable expression levels, the following analyses only included circRNAs supported by an average of at least five BSJ-spanning reads in each sample type. This resulted in 128 and 285 circRNAs (defined as high abundance circRNAs) in the lesional- and non-lesional skin, respectively. Most of these, 117 of 128 (91.4%) and 261 of 285 (91.6%), were also detected by the circExplorer bioinformatics algorithm that only considers circRNAs derived from annotated splice sites [29]. Forty-one (32.0%) and seventy-three (25.6%) circRNAs were on average expressed at higher levels than their linear host genes (circular-to-linear (CTL) ratio >1), and we detected four and two potentially novel circRNAs (not present in circBase [12]) in the lesional- and the non-lesional skin, respectively (Supplementary Table 2 and 3). Among the top 50 most abundant circRNAs in each sample type, we found four circRNAs present in both types that have previously been shown to be highly expressed in several different tissues in RNA-seq data from the ENCODE consortium (known as the top 10 alpha circRNAs (https://www.biorxiv.org/content/early/2018/05/25/328104) (Figure 1A and 1B). The 128 and 285 circRNAs found in our samples were generated from 115 and 241 different host genes, respectively (Figure 1C and 1D). Finally, we observed that circRNAs composed of two exons were most frequent in both sample types (Figure 1E and 1F).

**Figure 1.**
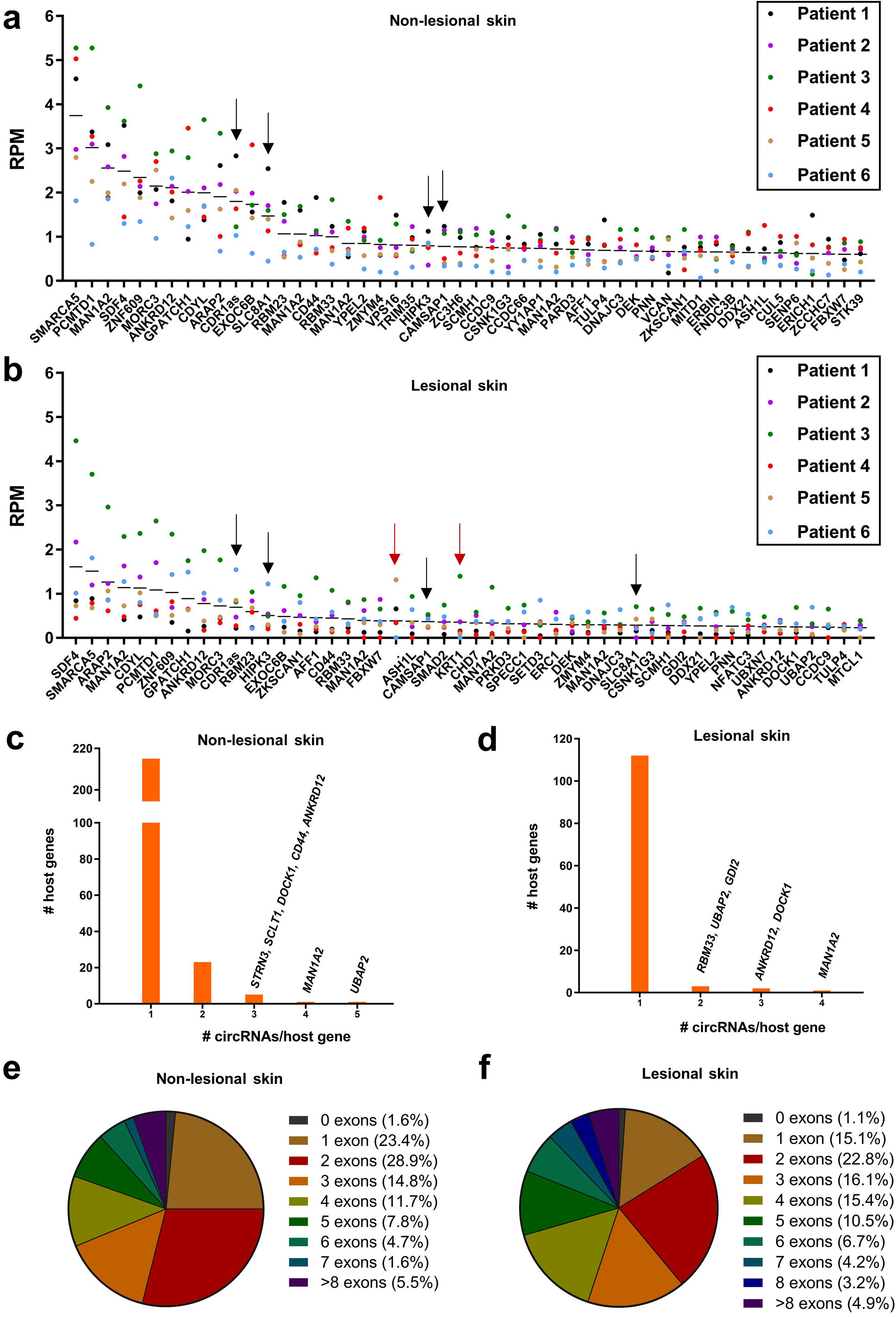
Characterization of circular RNAs in lesional- and non-lesional skin using RNA-seq. **(a-b)** Expression levels of the top 50 most abundant circRNAs in non-lesional skin **(a)** and lesional skin **(b).** Each dot represents the expression in reads per million (RPM) for a circRNA in one individual sample represented by a unique color. Each line represents the mean RPM. Black arrows indicate top 10 alpha circRNAs and red arrows indicated circRNAs not present in circBase. **(c-d)** Histograms showing the number of host genes producing various numbers of unique high abundance circRNAs in non-lesional skin **(c)** and in lesional skin **(d)**. **(e-f)** Pie charts showing the distribution of the numbers of exons annotated within the high abundance circRNAs in non-lesional skin **(e)** and lesional skin **(f)**.

### circRNAs are less abundant in lesional psoriatic skin relative to non-lesional skin

In total, we detected 298 unique high-abundance circRNAs in the lesional- and non-lesional skin biopsies combined. The overlap between circRNAs detected in the lesional- and non-lesional skin was considerable (38.6%), but many were unique mainly to the non-lesional skin (Figure 2A). In line with this, we observed that the 298 circRNAs were generally present at lower levels in lesional skin (Figure 2B). This was also true when comparing the lesional- and non-lesional skin for each individual patient (Supplementary Figure 1). In total, 148 circRNAs, including ciRS-7 (CDR1as), were significantly downregulated in lesional skin, while none was significantly upregulated (Figure 2C and Supplementary Table 4). To investigate if the dramatic downregulation of circRNAs in lesional skin was due to reduced expression of their host genes, fold changes in RPM were plotted against fold changes in CTL ratios. We observed that most of the circRNAs appeared along a diagonal line, indicating that they were downregulated independently of their host genes. However, there was also a substantial number of the downregulated circRNAs for which their respective host genes were downregulated to some extent (represented by dots appearing above the diagonal line in Figure 2D). In addition, we did not observe a considerable downregulation in lesional skin relative to non-lesional skin when considering other classes of RNA (Supplementary Figure 2).

**Figure 2.**
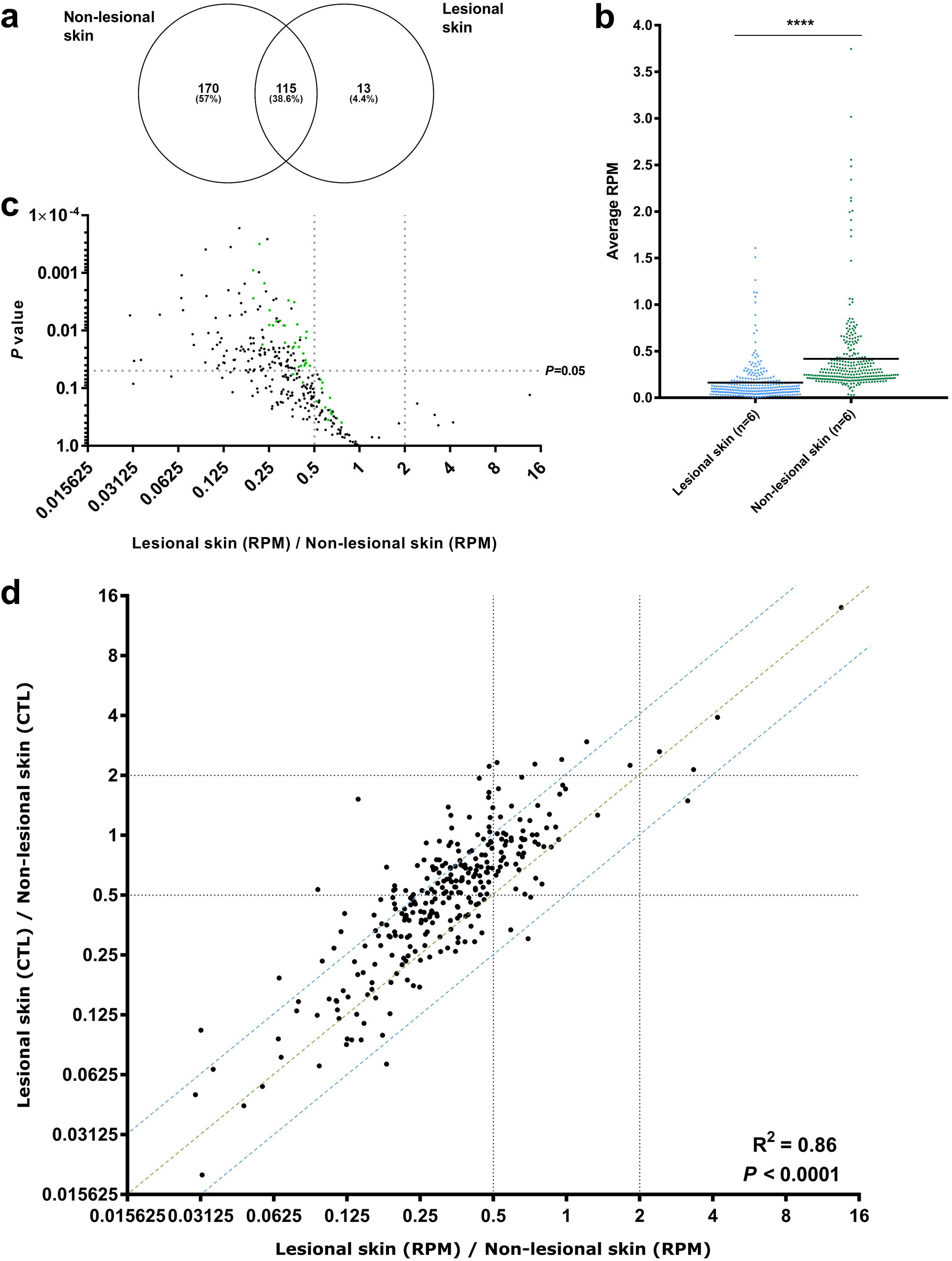
The circRNAome is massively downregulated in lesional- relative to non-lesional skin. **(a)** Venn diagram illustrating the overlap of circRNAs detected in the lesional- and the non-lesional skin. **(b)** Scatter plot showing that the average expression in reads per million (RPM) of the 298 unique high-abundance circRNAs is lower in lesional- relative to non-lesional skin. **(c)** Volcano plot of the 298 unique high-abundance circRNAs showing fold changes in circRNA expression in RPM between lesional- and non-lesional skin according to the levels of significance. The top 50 most abundant circRNAs are indicated in green. **(d)** Scatter plot of fold changes in RPM and fold changes in circular-to-linear (CTL) ratios of the high abundance circRNAs. It can be observed that most circRNAs were downregulated independent of their respective host genes (defined as those being present in between the blue lines).

Next, we wanted to rule out whether the observed downregulation was caused by experimental bias. Template switching and rolling circle amplification during reverse-transcription (RT), as well as PCR bias, are major concerns in the circRNA research field as artifacts mimicking circRNA molecules may arise because of these phenomena [35-37]. We therefore employed an enzyme-free technology (termed NanoString nCounter) [36] to profile the expression of the top 50 most abundant circRNAs in the entire dataset within the same samples. Again, we observed that circRNAs are dramatically downregulated in the lesional- relative to non-lesional skin (Supplementary Figure 3A). To extend the generality of the observed downregulation of circRNAs in psoriasis, we further analyzed six normal skin samples from unaffected individuals using our NanoString nCounter panel. Again, we observed a profound downregulation of circRNAs when comparing this data to data obtained from lesional skin (Supplementary Figure 3B). Finally, we analyzed a second cohort consisting of paired lesional- and non-lesional skin samples from another 13 psoriasis patients using our NanoString nCounter panel and confirmed the previous observations that circRNAs are generally less abundant in lesional skin (Supplementary Figure 3C).

### Factors known to influence circRNA biogenesis are unlikely to be the main drivers of circRNA downregulation in lesional skin

Several RNA-binding proteins, including *ADAR, FUS, DHX9, HNRNPL* and *QKI* have previously been shown to regulate circRNA biogenesis [16-20]. Thus, we speculated that expression changes of these genes might explain the observed downregulation of circRNAs in lesional skin.

Interestingly, *ADAR* was significantly upregulated in lesional skin both when considering the RNA-seq data (Figure 3A) and the NanoString nCounter data in both psoriasis cohorts (Figure 3B and Supplementary Figure 4). Because *ADAR* is known to suppress circRNA biogenesis mediated by base pairing between adjacent IAEs [16], we searched for IAEs within 2300 nucleotide regions flanking the BSJs (total distance) as previously described [16] for each of the high abundance circRNAs. The fraction of circRNAs with neighboring IAEs was 44% (131/298) and there was no tendency for circRNAs with IAEs to be more downregulated than circRNAs without them (*P*=0.75, chi-squared test) (Figure 3C). Consistent with this, the median distance to the nearest IAE did not differ significantly between circRNAs downregulated more than two-fold and the remaining circRNAs (*P*=0.10, Mann Whitney test). In addition, we did another search for IAEs within 10,000 nucleotide regions flanking the BSJs (total distance), this time requiring them to belong to the same subfamily as previously described [31]. There was no tendency for circRNAs with IAEs from the same subfamilies to be more downregulated than circRNAs without (*P*=0.60, chi-squared test) (Supplementary Figure 5A), and again the median distance to the nearest IAE did not differ significantly between circRNAs downregulated more than two fold and the remaining circRNAs (*P*=0.13, Mann Whitney test). Together, these analyses imply that other mechanisms are likely to be responsible for the observed downregulation of circRNAs in lesional skin.

**Figure 3.**
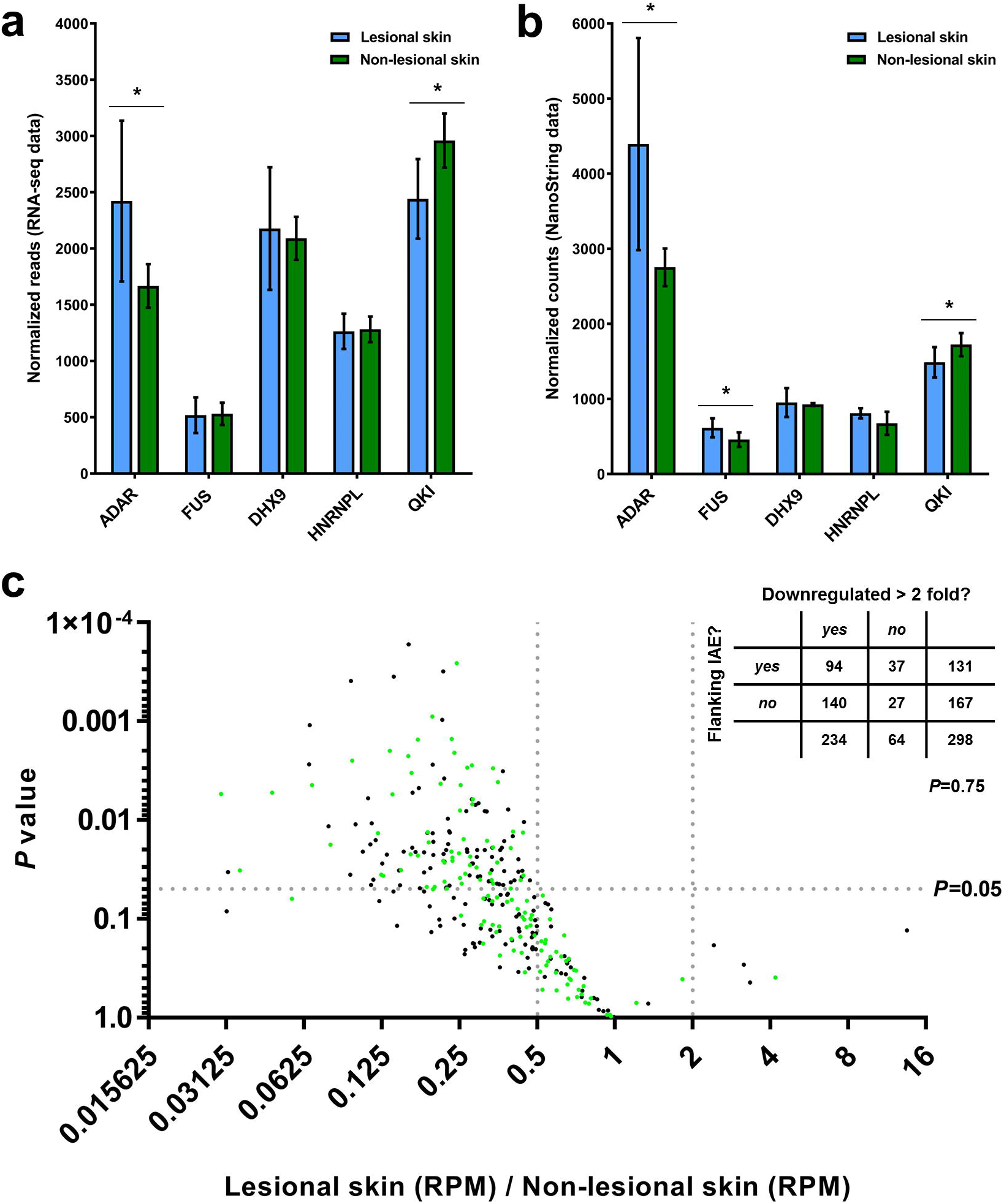
Analyses of factors known to regulate circRNA biogenesis in lesional- relative to non-lesional skin. **(a)** RNA-seq analyses revealed that *ADAR* was the most differentially expressed of the analyzed factors. **(b)** NanoString nCounter analysis confirmed that *ADAR* is upregulated in lesional- relative to non-lesional skin. **(c)** Volcano plot of the 298 unique high-abundance circRNAs showing fold changes in circRNA expression in RPM between lesional- and non-lesional skin according to the levels of significance. One-hundred thirty-one circRNAs likely to have *Alu*-mediated biogenesis are indicated in green (flanked by IAEs within 2300 nucleotide regions flanking the BSJs). Numbers of circRNA in each category are shown in the inserted table. There was no statistically significant association between downregulation of circRNAs (more than 2 fold) and the presence of flanking IAEs (*P* = 0.75, chi-squared test).

### circRNA expression in different cellular compartments of the skin biopsies

It is well known that lesional psoriatic skin contain more inflammatory cells such as lymphocytes than non-lesional skin [38, 39]. If the lymphocytes were to contain fewer circRNAs than keratinocytes, this could be a plausible explanation for the observed downregulation of the circRNAome in lesional skin. Before testing this hypothesis, we first confirmed that our lesional skin biopsies contained more lymphocytes than non-lesional skin. From hematoxylin and eosin staining of fixed skin biopsy sections, we found that especially the dermis of lesional skin showed greater lymphocytes numbers relative to the dermis of the non-lesional skin (Figure 4A). By analyzing the expression levels of T-cell specific genes [40], we could also confirm that the lesional skin contained more lymphocytes. In particular, four of six T-cell specific genes analyzed were significantly upregulated in the lesional skin (Figure 4B). Then, because the majority of the lymphocytes resides in the dermis, we separated the dermis from the epidermis by laser capture microdissection of five lesional skin- and five non-lesional skin biopsies and profiled the expression of the top 50 most abundant circRNAs using NanoString nCounter technology. Due to limited amounts of RNA, only seven circRNAs reached the detection limit in at least half of the samples. When comparing lesional and non-lesional epidermis, five of the seven circRNAs were significantly downregulated in the lesional skin, whilst the analysis of the dermis showed only one circRNA to be significantly different between the two tissue types (Figure 4C and 4D). In line with the above-mentioned analysis of factors known to affect circRNA biogenesis, we observed upregulation of *ADAR* in lesional skin. This applied to both the dermis and the epidermis (Figure 4E and 4F), supporting the notion that *ADAR* is not instrumental for the downregulation of the circRNAome in lesional skin.

**Figure 4.**
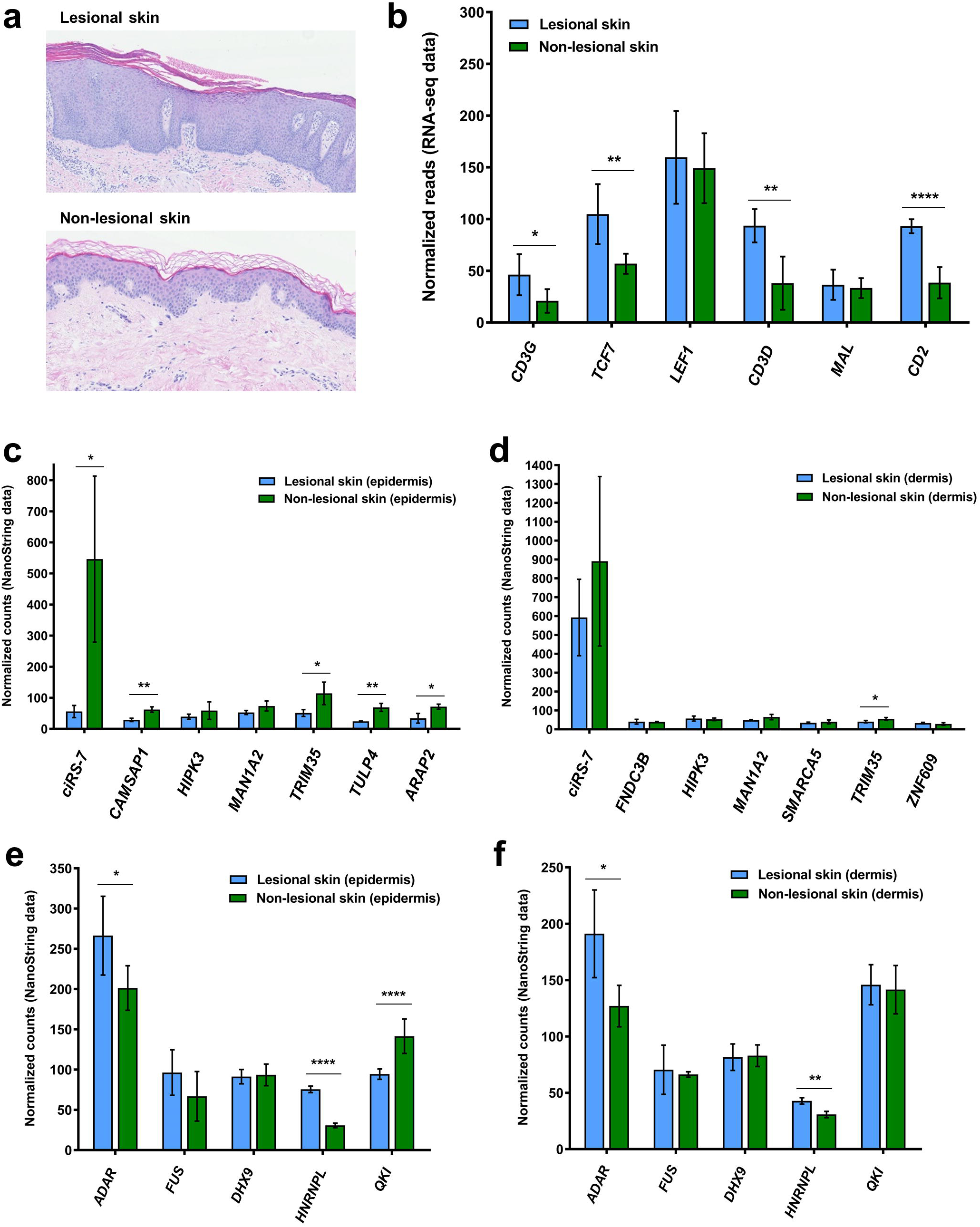
Downregulation of circRNAs is mainly observed in the epidermis and unlikely to be caused by infiltrating lymphocytes. **(a)** Representative H&E staining of lesional- and non-lesional skin samples showing a relative abundance of lymphocytes in the dermis of the lesional skin. **(b)** RNA-seq data showed that T-cell specific genes were expressed at higher levels in the lesional skin-relative to non-lesional skin. **(c-d)** Following microdissection of the epidermis and NanoString nCounter analysis, five of seven circRNAs were shown to be significantly downregulated in the epidermis of the lesional skin **(c)**, while this only applied to one of seven circRNAs analyzed in the dermis **(d)**. **(e-f)** *ADAR* proved to be significantly upregulated both in the epidermis **(e)** and in the dermis **(f)** of the lesional skin.

Next, to support our findings that circRNAs are mainly downregulated in the epidermis, we performed RNA chromogenic *in situ* hybridization (CISH) for ciRS-7, since this circRNA was significantly downregulated in both the RNA-seq data and in the NanoString nCounter data for both cohorts. Consistent with these analyses, we observed a marked downregulation of ciRS-7 in the epidermis of lesional- vs non-lesional skin (Figure 5), while ciRS-7 expression in the dermis was comparable between lesional and non-lesional skin.

**Figure 5.**
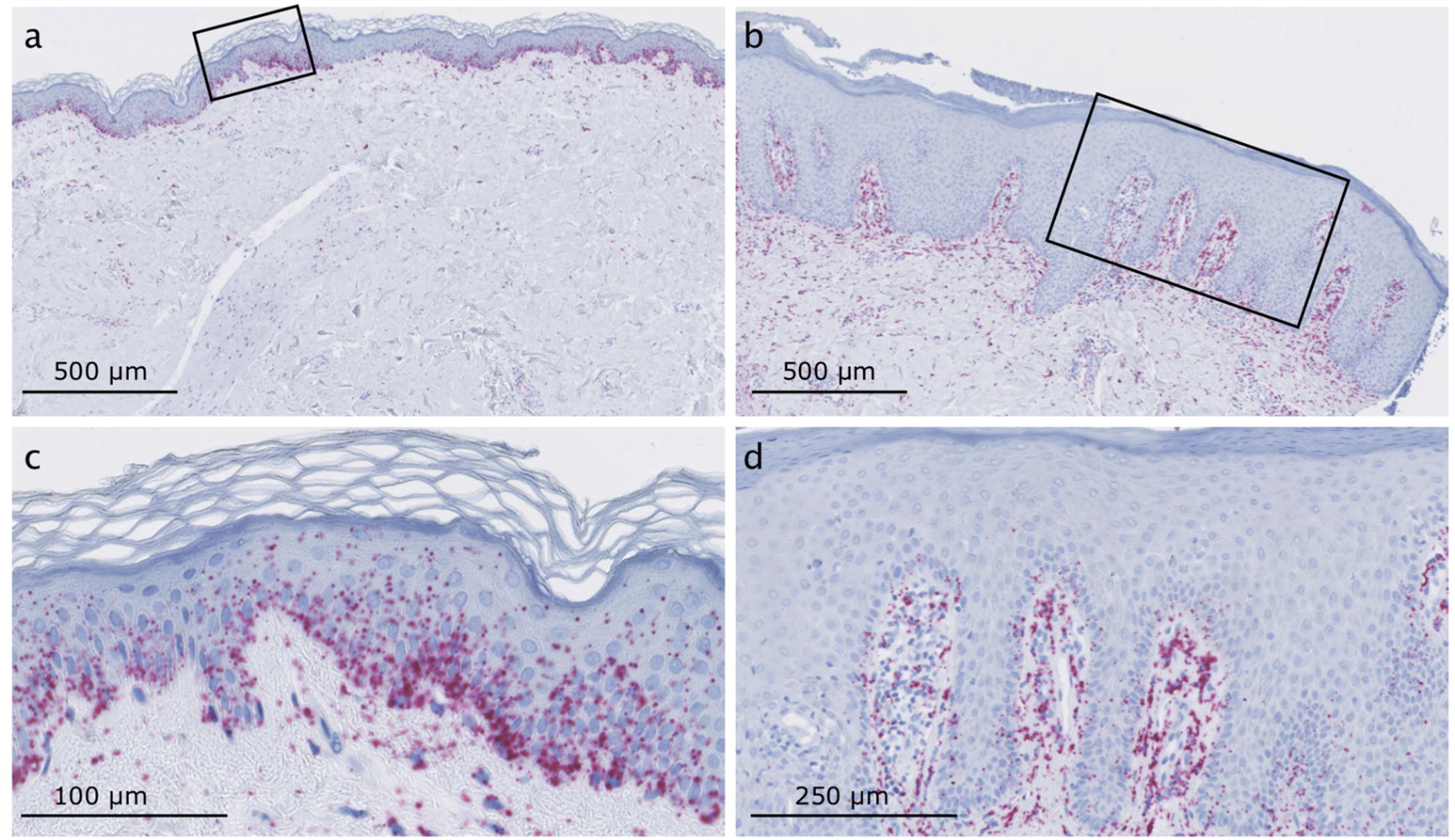
RNA chromogenic *in situ* hybridization (CISH) for ciRS-7 in lesional- and non-lesional skin. **(a-b)** Overviews with the areas shown in **(c-d)** indicated with a square. The ciRS-7 signal (red dots) is observed mainly in the epidermis of the non-lesional skin **(a and c)** and in the dermis of lesional skin **(b and d)**. Scale bars are indicated in the lower right corner.

### Identification and characterization of miRNAs in lesional- and non-lesional skin biopsies

Having established that circRNAs are profoundly downregulated in lesional skin, we speculated that this phenomenon could be responsible for previously reported alterations in miRNA expression associated with psoriasis [4, 5]. To investigate this, we profiled the expression of approximately 800 miRNAs using the nCounter Human v3 miRNA panel from NanoString Technologies, in the lesional and non-lesional skin biopsies.

We detected 182 and 143 unique miRNAs supported by at least two counts in a single sample in the lesional (n=6) and non-lesional skin biopsies (n=6), respectively. In order to ensure that we look at miRNAs with reasonable expression levels, we only considered miRNAs supported by an average of at least five counts (defined as high abundance miRNAs) in each sample type. This resulted in a total of 123 and 106 miRNAs in the lesional skin- and non-lesional skin biopsies, respectively.

Combining the lesional- and non-lesional skin samples, we detected 137 unique high abundance miRNAs, with a substantial overlap between the miRNAs detected in each sample type (67.2%) (Figure 6A). When considering all patients combined, the median expression of the high abundance miRNAs were significantly higher in lesional- relative to non-lesional skin (Figure 6B). However, this did not apply to each individual patient (Supplementary Figure 6). In total, 37 miRNAs were differentially expressed; 31 were upregulated in the lesional skin and six were downregulated (Supplementary Table 5). Several of these miRNAs have been shown to be differentially expressed in psoriatic skin in previous studies and are highlighted in Figure 6C.

**Figure 6.**
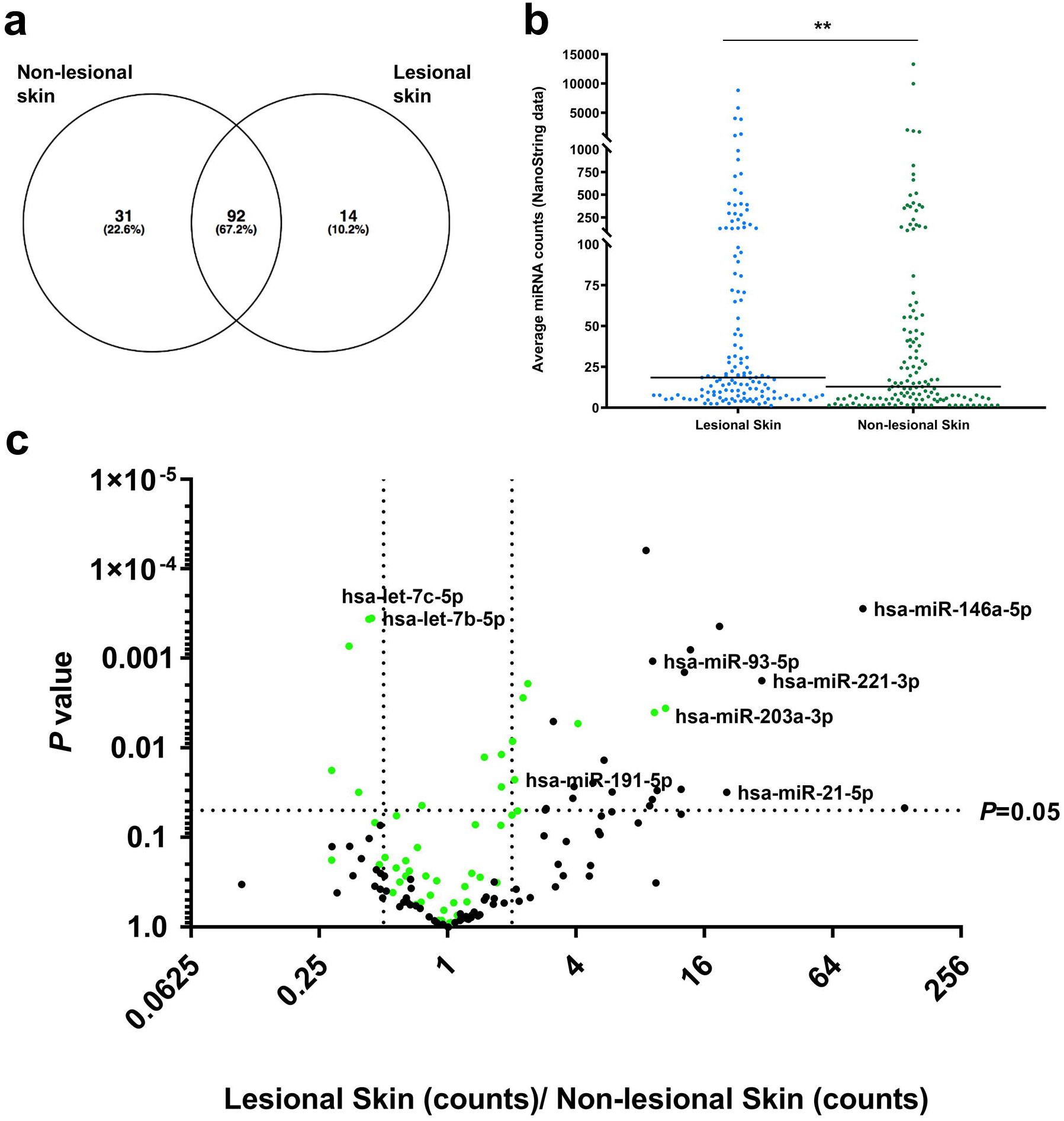
Characterization of miRNAs in lesional skin- and non-lesional skin using NanoString nCounter analysis. **(a)** Venn diagram illustrating the overlap between the miRNAs detected in lesional- and non-lesional skin. **(b)** The average expression of the 137 high abundance miRNAs is slightly lower in lesional- relative to non-lesional skin. **(c)** Volcano plot of the 137 high abundance miRNAs with the top 50 most abundant miRNAs indicated in green and names of miRNAs previously shown to be differentially expressed in psoriasis indicated.

### Investigation of potential interactions between circRNAs and miRNAs

To analyze potential interactions between miRNAs and circRNAs in psoriasis, we considered only the high abundance circRNAs and miRNAs as the quantitative data for these are much more reliable than for circRNAs and miRNAs supported by very few reads. We first predicted miRNA binding sites in each of the individual circRNAs and multiplied the number of sites with the change in expression (ΔRPM values) between lesional- and non-lesional skin for each corresponding circRNA. The sum of the ΔRPM values for each miRNA binding site was plotted against the fold changes (Figure 7A) and absolute changes (Figure 7B) in miRNA expression levels between lesional- and non-lesional skin, respectively. For both of these analyses, we did not observe any significant correlations between the loss of miRNA binding sites on the circRNAs and differences in miRNA expression levels between lesional and non-lesional skin. The best candidate for a miRNA that was differentially expressed as a result of changes in the circRNAome was miR-203a-3p (Figure 7) as many miR-203a-3p binding sites were present in downregulated circRNAs and miR-203a-3p was among the most upregulated miRNAs in lesional skin. However, this does not seem to reflect a general interplay between circRNAs and miRNAs in our samples.

**Figure 7.**
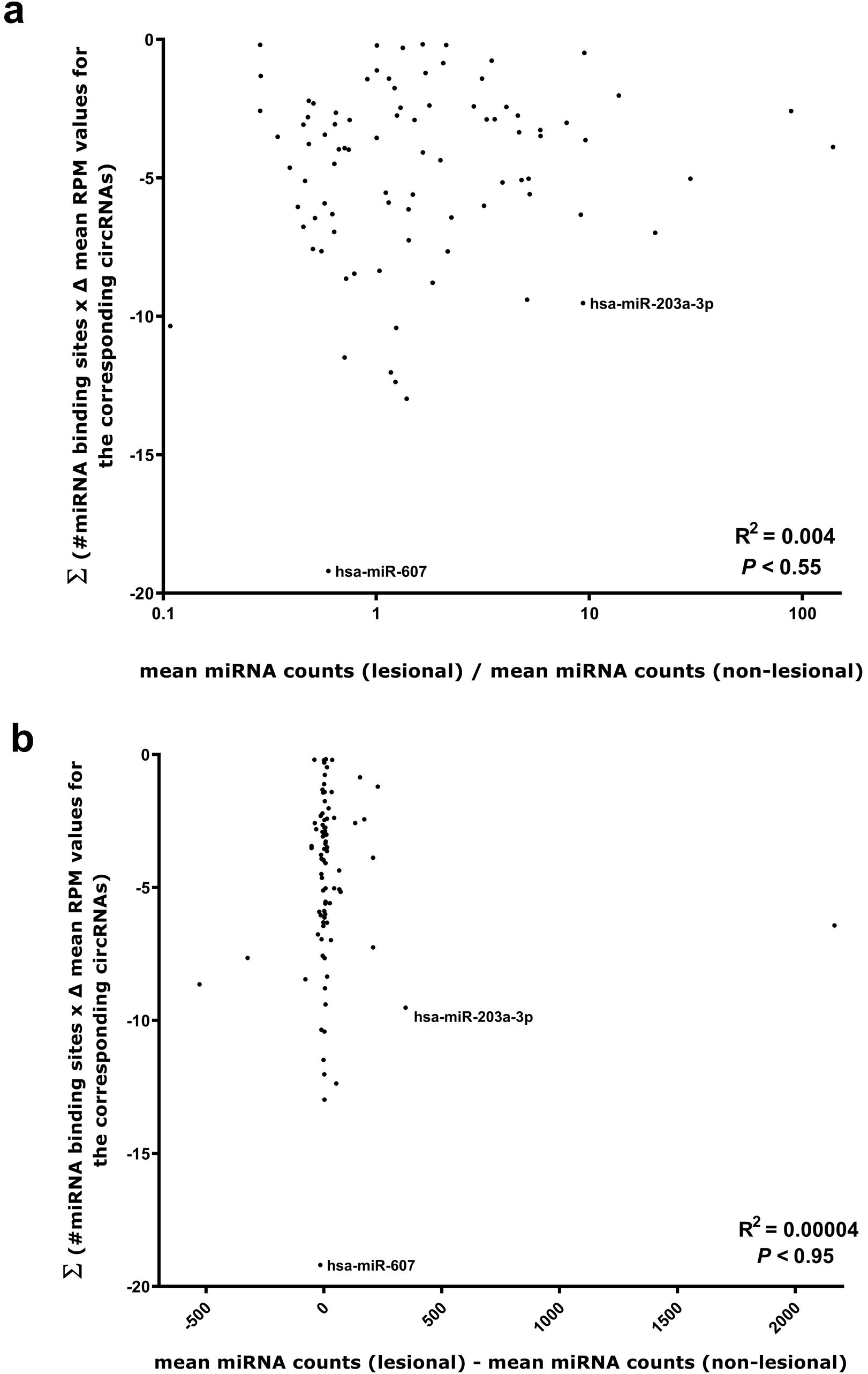
Changes in the amounts of available miRNA binding sites present on circRNAs do not correlate with changes in miRNA expression levels in lesional- relative to non-lesional skin. **(a-b)** For each miRNA, the sum of the number of binding sites for that particular miRNA was multiplied with the mean circRNA RPM (lesional) -mean circRNA RPM (non-lesional) for the circRNAs harboring binding sites for that particular miRNA. These values were plotted against either the fold change (mean miRNA counts (lesional)/mean miRNA counts (non-lesional)) for each high abundance miRNA with at least one binding site present on the high abundance circRNAs **(a)** or the absolute difference in expression level (mean miRNA counts (lesional) -mean miRNA counts (non-lesional)) for each high abundance miRNA with at least one binding site present on the high abundance circRNAs **(b)**.

## Discussion

In this study, we present the first sequencing-based catalogue of circRNA expression in lesional- and non-lesional skin from patients diagnosed with psoriasis vulgaris. Our data show that the circRNAome is extensively downregulated in lesional skin. The RNA-seq data were confirmed using an enzyme-free method [36], which is not subject to PCR bias and potential artifacts related to reverse transcription [35-37], and could be extended to another patient cohort. In addition, we also observed an overall downregulation of circRNAs when comparing to normal healthy skin.

The circRNAome was mainly downregulated in the epidermis of the skin, an effect that could not be explained by differential expression of known circRNA biogenesis factors nor by differences in the number of infiltrating lymphocytes. We did observe that *ADAR* was expressed at significantly higher levels in the lesional skin but there was no tendency for circRNAs with IAEs in the introns flanking their BSJs to be more downregulated than circRNAs without, which argues against upregulation of *ADAR* as the main mechanism responsible for downregulation of the circRNAome in psoriasis. In addition, despite being statistically significant, the difference in *ADAR* expression was less than two fold. Thus, a more likely explanation for the altered circRNA levels may involve the higher proliferation and turnover rates of the keratinocytes in lesional skin, and the fact that those keratinocytes are less likely to become terminally differentiated. It has previously been observed that circRNAs are upregulated during differentiation [15, 22] and that higher levels of these molecules are found in non-proliferating cells [41]. In particular, circRNA molecules have very long half-lives in cells due to their slow turnover rates [9] and the relative inefficiency of the back-splicing reaction required to form most circRNAs imply that they generally need longer time to build up to steady-state levels. Thus, we propose that the less differentiated and highly proliferative keratinocytes in the lesional psoriatic skin prevents the highly stable circRNA molecules from building up to the same expression levels found in the non-lesional skin. In support of this hypothesis, we observed that most of the circRNA expression changes were independent of expression changes of their respective linear host genes. In other words, less active promoters and enhancers may only explain a minor part of the observed downregulation of the circRNAome. In addition, when analyzing different classes of linear RNA, including miRNAs, mRNAs, lincRNAs, antisense RNAs, and snoRNAs, we did not observe an overall downregulation in lesional skin relative to non-lesional skin.

Interestingly, a recent study has shown that circRNAs are downregulated as a class upon viral infection [42]. Using genome-wide siRNA screening targeting all human unique genes and an efficient circRNA expression reporter, this study identified the immune factors NF90/NF110 as key regulators in circRNA biogenesis. The gene encoding these factors (*ILF3*) was not differentially expressed in our RNA-seq data, which was expected since the immune pathways activated in psoriasis are different from those activated upon viral infection. However, we cannot exclude that other yet unknown factors could play a role in the downregulation of circRNAs in psoriasis.

Deregulation of miRNA levels has been implicated in the pathogenesis of psoriasis and numerous studies have indicated that circRNAs may function as efficient inhibitors of miRNA activity [10, 21, 26, 43]. Therefore, we hypothesized that loss of miRNA binding sites, as a consequences of the downregulated circRNAome, could explain some of the changes in miRNA expression previously observed in psoriasis. To test this hypothesis, we profiled the expression of a large number of miRNAs in the skin biopsies using the nCounter Human v3 miRNA panel from NanoString Technologies, and quantified the number of individual miRNA binding sites in each high abundance circRNA. However, we were unable to find any correlations between the loss of miRNA binding sites on the circRNAs and differences in miRNA expression levels between lesional and non-lesional skin. Together, our observations suggest that the global reduction in circRNA expression in lesional skin is more likely to be a consequence than a cause of the disease. However, this does not necessarily imply that individual circRNAs may not be directly involved in psoriasis pathogenesis.

In fact, a recent study argues that circRNAs are directly involved in the pathogenesis of psoriasis [44]. The authors of this study performed RNA-seq of stem cells isolated from lesional psoriatic skin and normal healthy skin. However, the patients had a 1-month to 20-year disease course and a wide range in disease severity (PASI scores between 3.0 and 43.8), and the stem cells were cultured for five generations before RNA-seq analyses. The bioinformatics algorithm CIRI was used for circRNA quantification, which we have previously shown to have a low accuracy for circRNA detection [45], and indeed more than half of the 6,000 circRNAs detected was not present in circBase. The authors did not validate the circular nature of any of these potentially novel circRNAs by assessing their resistance to degradation by RNase R and no comparisons to other bioinformatics algorithms were performed [44]. In contrast to our study, the authors found many more upregulated-than downregulated circRNAs and went on to functionally study a potentially novel circRNA derived from chr2:206992521|206994966. This circRNA was predicted to interact with several miRNAs, which were predicted to regulate STAT3, STAT4 and IL2RB, however, no experimental evidence was provided in support of this [44]. We did not find this particular circRNA among the more than 3,000 unique circRNAs detected in the lesional- and non-lesional skin biopsies and could, therefore, not examine if it is differentially expressed in our samples.

Another recent study profiled circRNA expression in three psoriatic lesions and three normal healthy skin tissues [46]. The RNA from these samples was amplified prior to microarray analysis. The authors found almost 5,000 differentially expressed circRNAs of which the majority was upregulated in psoriasis. Six differentially expressed circRNAs were then selected for verification by RT-qPCR, but only one of these, hsa_circ_0061012, could be verified [46]. Again, we did not detect this circRNA and could not examine if it is differentially expressed in our samples.

To the best of our knowledge, our study is the first to use the NanoString nCounter technology to profile miRNA expression in psoriasis and, in accordance with earlier reports, we found that miR-146a, miR-21, miR-203, miR-221, miR-155 and miR-223 were significantly upregulated, and that hsa-let-7b-5p, hsa-let-7c-5p and miR-125b were significantly downregulated in lesional- relative to non-lesional skin [4, 5]. Additionally, we identified several miRNAs that have not previously been associated with psoriasis. For instance, miR-200c-3p, known to enhance the invasive capacity of human squamous cell carcinoma [47], miR-191-5p, abnormally expressed in several cancers and various other diseases [48], and hsa-miR-93-5p, promoting cancer cell proliferation through inhibiting *LKB1* in lung adenocarcinoma [49], were all significantly upregulated in lesional skin.

## Conclusions

In this study, we have found a global reduction of circRNA expression levels in lesional- relative to non-lesional skin from patients diagnosed with psoriasis. This phenomenon was mainly restricted to the epidermis and could not be explained by expression changes in factors known to affect circRNA biogenesis nor by differences in the number of lymphocytes in the samples. Instead, we suggest that the downregulated circRNAome in lesional skin may be related to a passive dilution of the circRNAs caused by the high proliferation- and turnover rates of the keratinocytes in the epidermis of the skin. It is too early to say whether the altered circRNA expression in psoriasis is a cause or consequence of the disease, however, our data do not support an active role for circRNAs in psoriasis pathogenesis via alteration of miRNA expression levels.

## Declarations

## Acknowledgements

We thank Birgit MacDonald, Malene Hykkelbjerg Nielsen and Birgit Roed Sørensen for their outstanding technical assistance and Anne Færch Nielsen for a thorough and critical reading of the manuscript.

## Funding

This project has received funding from the European Union’s Horizon 2020 research and innovation programme under the Marie Skłodowska-Curie grant agreement No 721890, and LSK is supported by a grant from the Carlsberg Foundation (CF16-0087).

## Availability of data and materials

The datasets supporting the conclusions of this article are included within the article and its supplementary files. Our raw data cannot be submitted to publicly available databases because the patients were not consented for sharing their raw data, which can potentially identify the individuals, but are available from the corresponding author on reasonable request.

## Authors’ contributions

LSK, LM, LI, CJ and JK conceived the study. CJ and LI collected the clinical samples used. LM and LSK carried out the laboratory work related to RNA-seq and NanoString nCounter analyses. TLA carried out the RNA CISH analysis. HH carried out microdissections and H&E stainings. TLHO and TBH carried out the bioinformatics analysis related to IAE in BSJ flanking regions. MTV and LSK performed the bioinformatics analyses related to RNA-seq experiments. LM and LSK analyzed all data and prepared the figures and tables. JK and LSK provided reagents and materials. LSK wrote the final version of the manuscript, which was revised for important intellectual content and approved by all authors.

## Consent for publication

Not applicable.

## Competing interests

The authors declare no competing financial interests.

## Supplementary figure legends

**Supplementary Figure 1. (a-f)** Scatter plots showing the average expression in reads per million (RPM) of the 298 unique high-abundance circRNAs in lesional- and non-lesional skin for each of the individual patients. The bars represent means.

**Supplementary Figure 2. (a)** Volcano plot of the top 298 most abundant mRNAs showing fold changes in mRNA expression between lesional- and non-lesional skin according to the levels of significance. Several genes know to be upregulated in psoriasis are indicated. **(b)** Volcano plot of the top 298 most abundant lincRNAs showing fold changes in lincRNA expression between lesional- and non-lesional skin according to the levels of significance. **(c)** Volcano plot of the top 298 most abundant antisense RNAs showing fold changes in antisense RNA expression between lesional- and non-lesional skin according to the levels of significance. **(d)** Volcano plot of the top 298 most abundant snoRNAs showing fold changes in snoRNA expression between lesional- and non-lesional skin according to the levels of significance.

**Supplementary Figure 3. (a)** Volcano plot of the top 50 most abundant circRNAs showing fold changes in circRNA expression in counts between lesional- and non-lesional skin according to the levels of significance. **(b)** Volcano plot of the top 50 most abundant circRNAs showing fold changes in circRNA expression in counts between lesional skin and healthy control skin according to the levels of significance. **(c)** Volcano plot of the top 50 most abundant circRNAs showing fold changes in circRNA expression in counts between lesional- and non-lesional skin from a second cohort of 13 patients according to the levels of significance. All analyses were performed using our custom NanoString nCounter panel.

**Supplementary Figure 4.** NanoString nCounter analysis confirmed that *ADAR* is upregulated in lesional- relative to non-lesional skin samples from a second cohort of 13 patients.

**Supplementary Figure 5.** Volcano plot of the 298 unique high-abundance circRNAs showing fold changes in circRNA expression in RPM between lesional- and non-lesional skin according to the levels of significance. Two-hundred thirty-three circRNAs likely to have *Alu*-mediated biogenesis are indicated in green (flanked by IAEs from the same subfamily within 10,000 nucleotide regions flanking the BSJs). Numbers of circRNA in each category are shown in the inserted table. There was no statistically significant association between downregulation of circRNAs (more than 2 fold) and the presence of flanking IAEs from the same subfamily (*P* = 0.60, chi-squared test).

**Supplementary Figure 6. (a-f)** Scatter plots showing the average expression in counts of the 137 unique high-abundance miRNAs in lesional- and non-lesional skin for each of the individual patients. The bars represent means.

