## Supplementary figures and images for "High-throughput RNA sequencing from paired lesional- and non-lesional skin reveals major alterations in the psoriasis circRNAome"

### Supplementary Figure 1

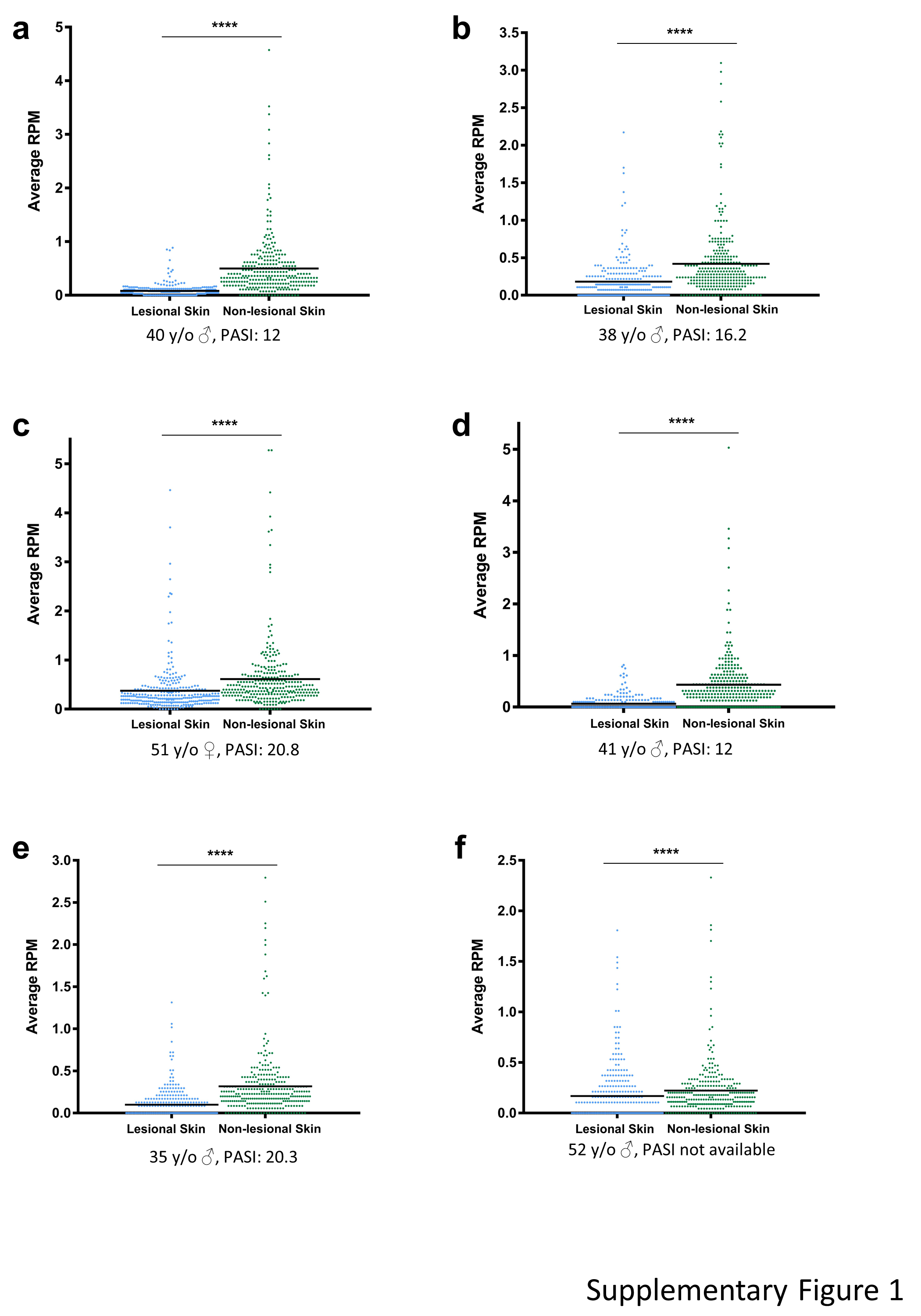

### Supplementary Figure 2

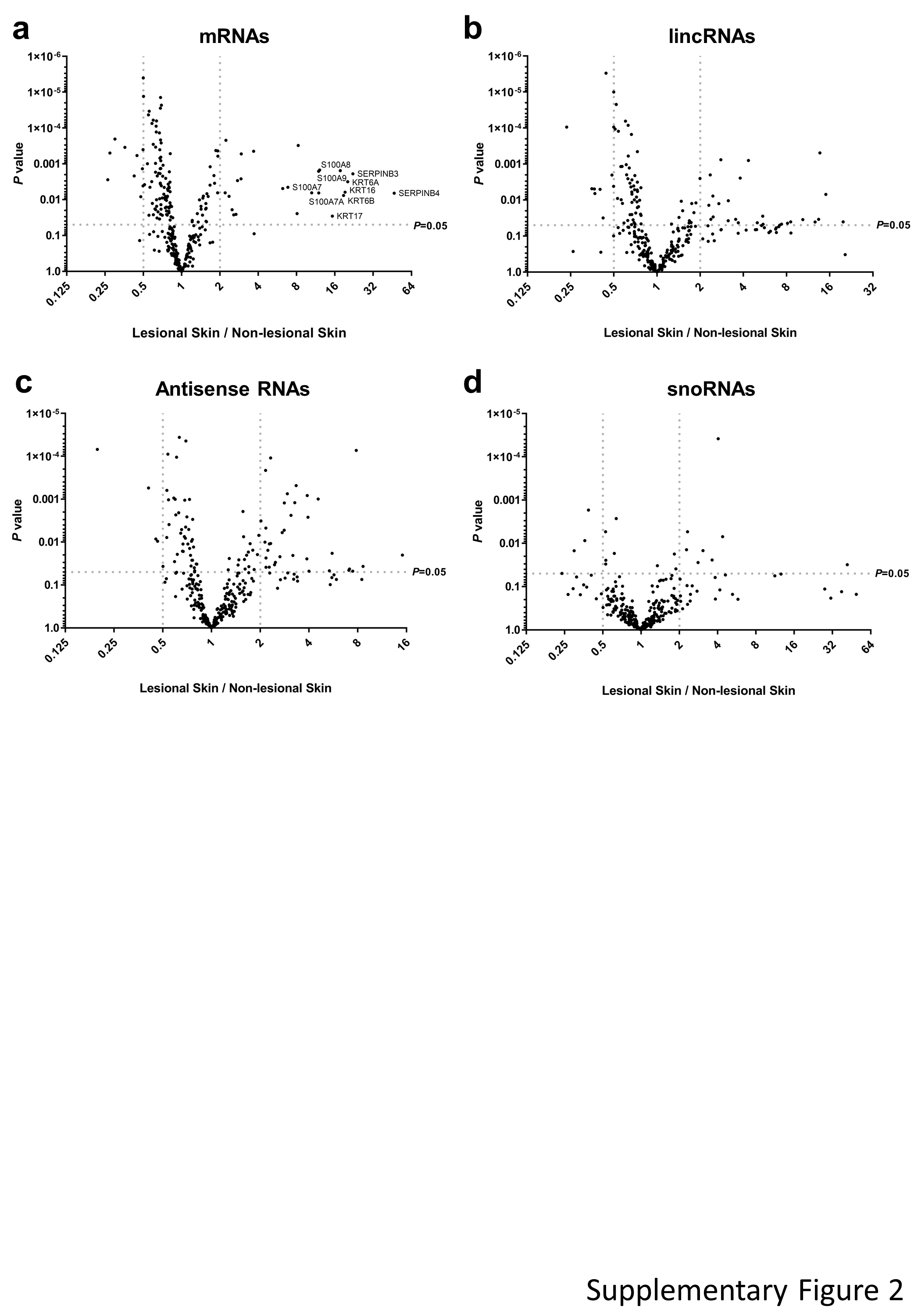

### Supplementary Figure 3

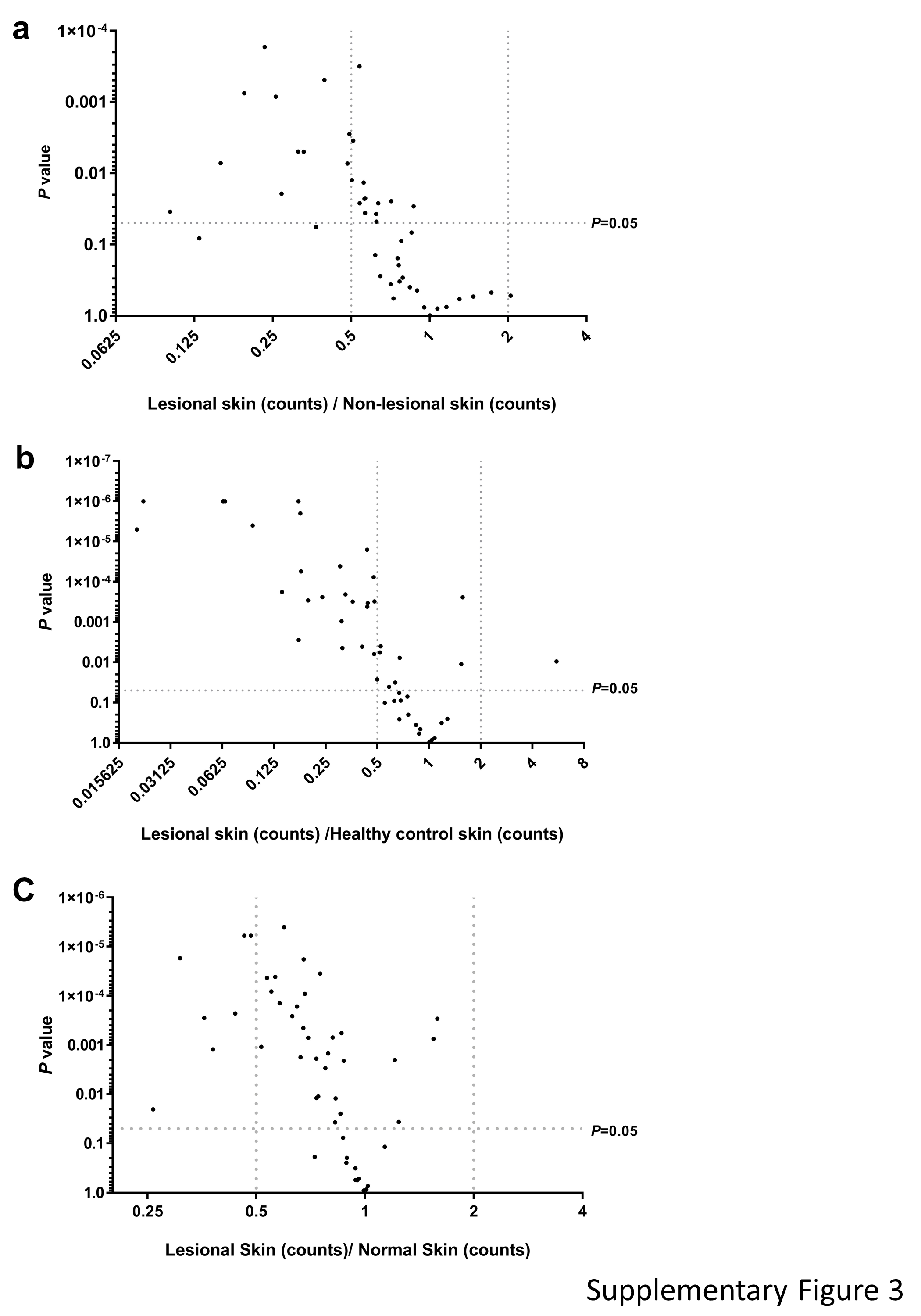

### Supplementary Figure 4

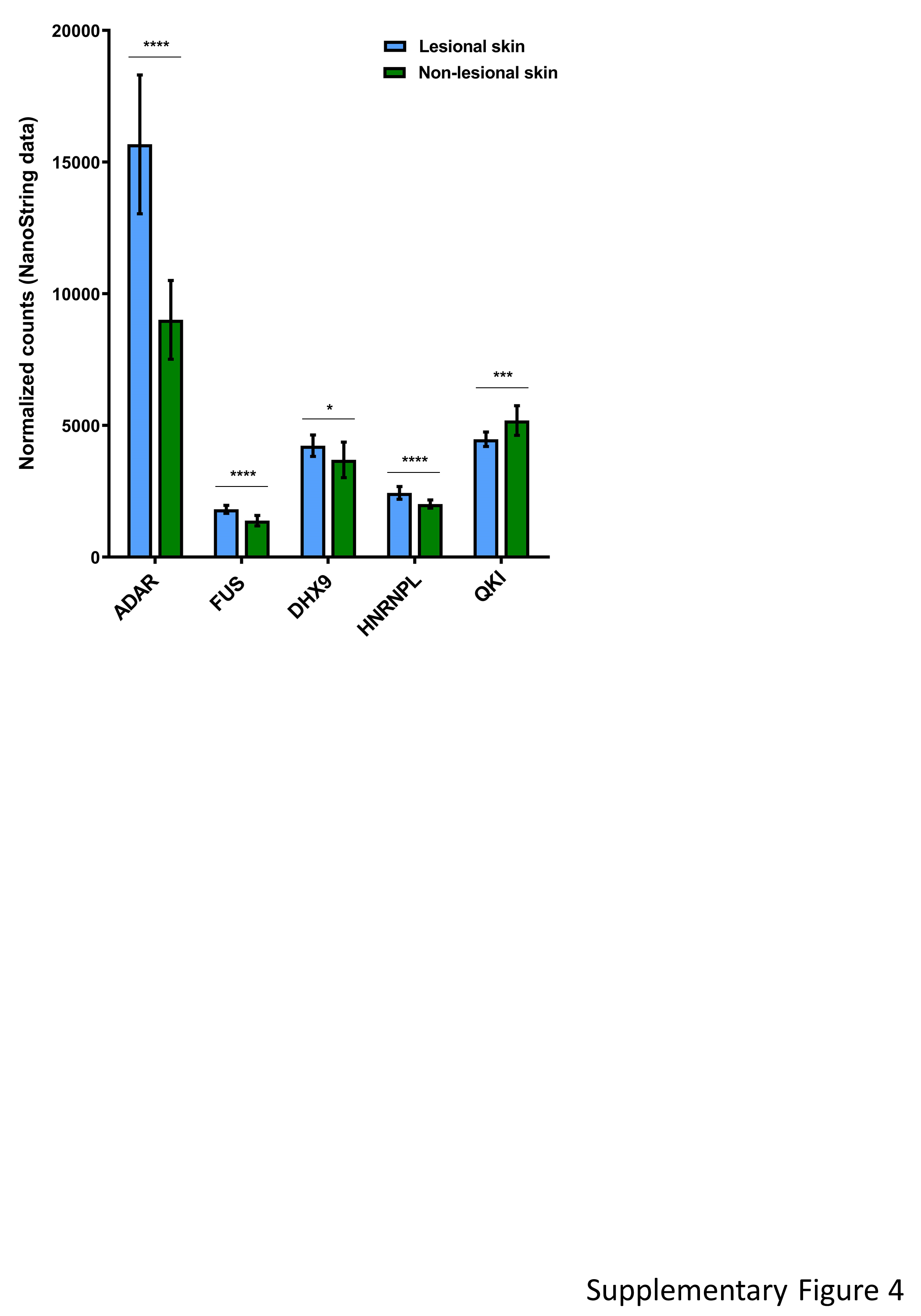

### Supplementary Figure 5

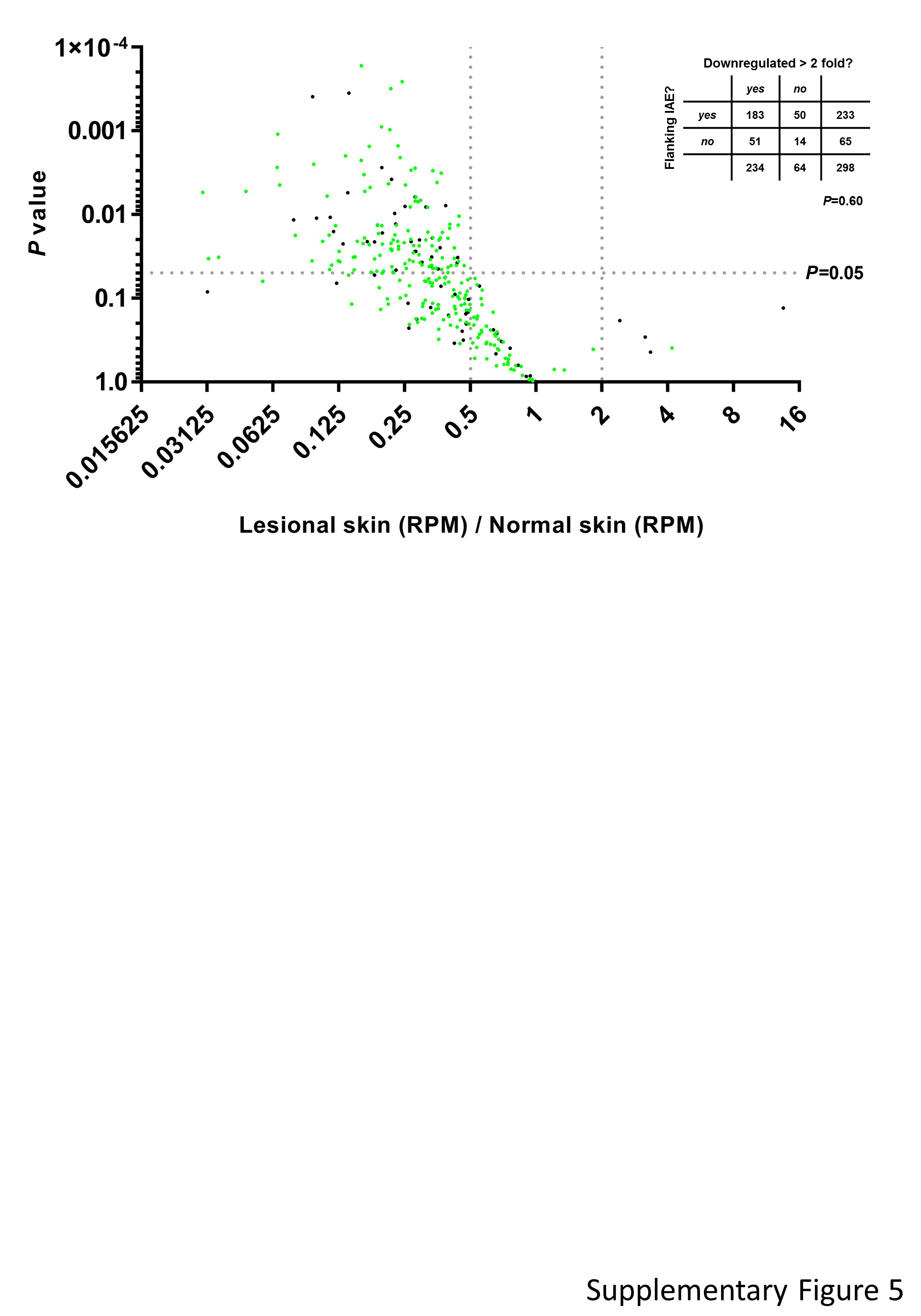

### Supplementary Figure 6

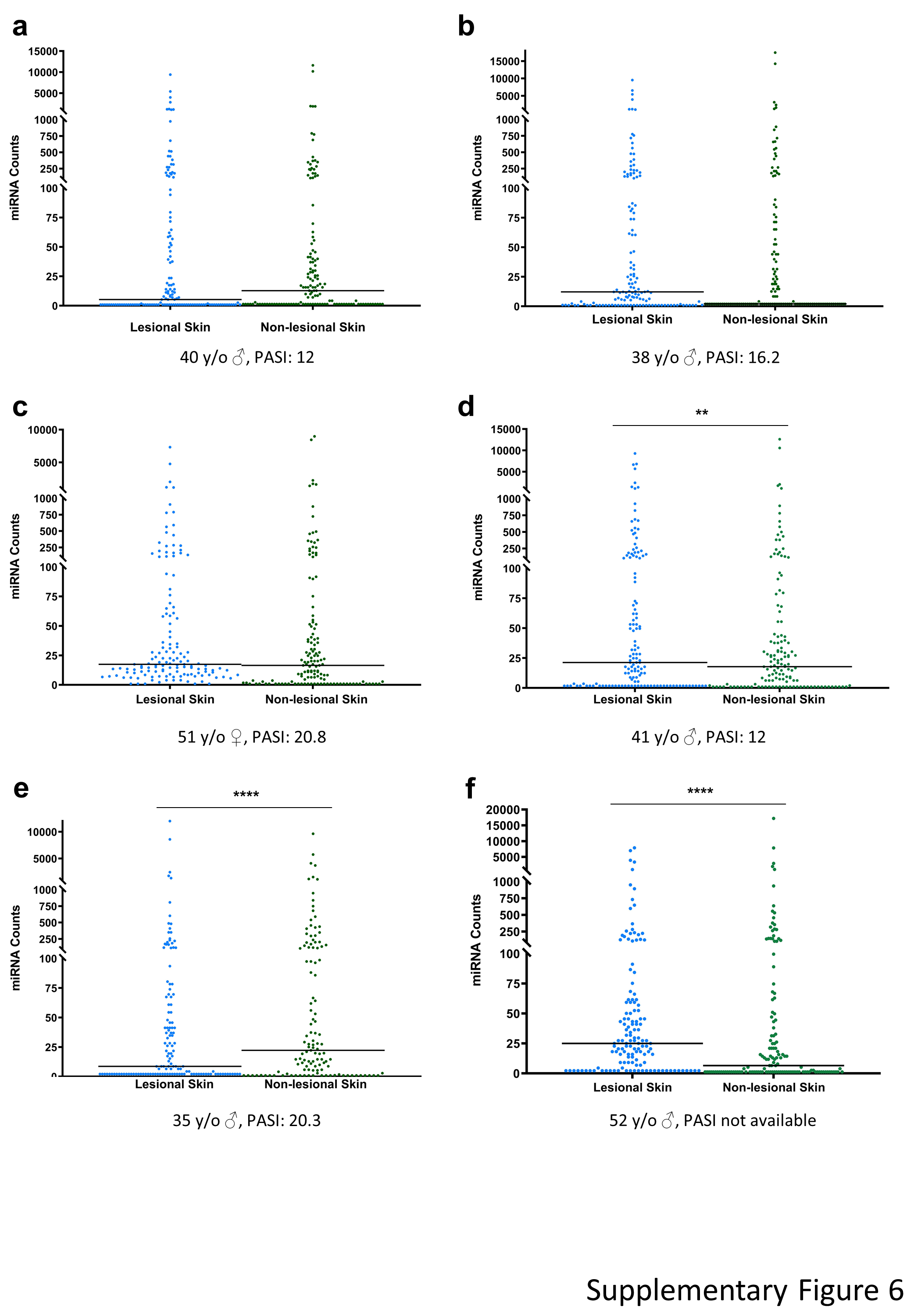
