## Supplementary Tables for "High-throughput RNA sequencing from paired lesional- and non-lesional skin reveals major alterations in the psoriasis circRNAome"

**Supplementary Table 1.** Custom CodeSet of capture- and reporter probes designed to target regions of 100 bp overlaying the BSJs of the top 50 most abundant circRNAs in the entire dataset.

| **Class** | **Name** | **Target sequences** |
| --- | --- | --- |
| mRNA | ACTB | TGCAGAAGGAGATCACTGCCCTGGCACCCAGCACAATGAAGATCAAGATCATTGCTCCTCCTGAGCGCAAGTACTCCGTGTGGATCGGCGGCTCCATCCT |
| mRNA | ADAR | TATGCCCAGTTCGCTAGTCAAACCTGTGAGTTCAACATGATAGAGCAGAGTGGACCACCCCATGAACCTCGATTTAAATTCCAGGTTGTCATCAATGGCC |
| circRNA | AFF1 | CTGCCAAAGCCAAGCTCACCAAACTGAAGATGCCTTCTCAGTCAGTTGAGTTTGTACAATGACGACAGAAACCTGCTTCGAATTAGAGAGAAGGAAAGAC |
| circRNA | ALAS1 | AGAAAGCAGGCAAATCTCTGTTGTTCTATGCCCAAAACTGCCCCAAGATGATGGAAGTTGGGGCCAAGCCAGCCCCTCGGGCATTGTCCACTGCAGCAGT |
| circRNA | ANKRD12 | GCTACTAAAAAGAGAGGTGCCTTTATCTGATGATGATGAAAGTTACACAGATCCAGGATGAGAAGACTGATAAAAGAAGAAGCTAGCTGAACAGCTGTAA |
| circRNA | ARAP2 | CATCGACCTGTACCAGAGATTCCAGGGTCAACAAAAGGAGTTTCTGGGAGGCTTAAGCAGGTAAAGCAGTCACTGTGAAGAAAATAACATTTTAAGAAAC |
| circRNA | ASH1L | AAAAAGCAGAAGCCATTACCAGAGGAAGAAGAGCAAGAGAATAATAAAAGGCTCCCAGCCAACCTCTGATAAACCCTCCCAGCGGCCATCAGAGAGCACA |
| mRNA | B2M | CGGGCATTCCTGAAGCTGACAGCATTCGGGCCGAGATGTCTCGCTCCGTGGCCTTAGCTGTGCTCGCGCTACTCTCTCTTTCTGGCCTGGAGGCTATCCA |
| circRNA | CAMSAP1 | AAGAACTTCCGTACGACCTCGAGGATGCCATGGTGTTCTGGATCAACAAGATAACATCCCTGAGGACCTCAGAGACCCTTTCTACGTTGACCAGTATGAG |
| circRNA | CCDC66 | CTCAGATTGAGGAACGAGACAGACGACGACAAAAACAATTAGAGCATCAGGAAACAGTACTGCTGGAGCACCCTTTCAGTGCTGTGAAACAAGAACTGCA |
| circRNA | CCDC9 | GGGGCCCCGACTTCGAGCGGGTGCGCTGTGGCCTTGAGCACGAGCGGCAGGAGTGGGAGGAGCGGCGCAGGCAGAACATTGAGAAGATGAATGAGGAGAT |
| circRNA | CD44 | ATATCGCCAAACACCCAAAGAAGACTCCCATTCGACAACAGGGACAGCTGTTTCAACCACACCACGGGCTTTTGACCACACAAAACAGAACCAGGACTGG |
| circRNA | CDR1 | AACGTCTCCAGTGTGCTGATCTTCTGACATTCAGGTCTTCCAGTGTCTGCAATATCCAGGGTTTCCGATGGCACCTGTGTCAAGGTCTTCCAACAACTCC |
| circRNA | CDYL | CGGCCCTGTGACTGCAGCCATGGCCACAGGCTTAGCTGTTAACGGGAAAGGTTGAAAGGATTGTTGACAAAAGGAAAAATAAAAAAGGGAAGACAGAGTA |
| circRNA | CSNK1G3 | AAATCAAGAGCACCACAGCTACATTTGGAATACAGATTCTATAAGCAGTTAGGATCTGGAGCTCTCTATCAATATCAGCTCACATCATTGAAAAGATAAT |
| circRNA | CUL5 | GAGCACTACGTTATTTAGAAACAAGACGAGAATGTAACTCCGTTGAAGCAGGATGTGCATGCAGTCTGTCTTTGGGATGATAAAGGCCCAGCAAAAATTC |
| circRNA | DDX21 | GTAACCCCAGTGAAGCTGCCAGTGAAGAAAGTAACAGTGAGATAGAGCAGAAAGAGAAAAAAGAGAAGCCAAAATCTGATAAGACTGAAGAGATAGCAGA |
| circRNA | DEK | TGGCCAGTGCTAACTTGGAAGAAGTCACAATGAAACAGATTTGCAAAAAGAAAAGAGTCTCATCGTGGAAGGCAAGAGGGAAAAGAAAAAAGTAGAGAGG |
| mRNA | DHX9 | AATGAACGTATGCTGAACATGATCCGTCAGATCTCTAGACCCTCAGCTGCTGGTATCAACCTTATGATTGGCAGTACACGGTATGGAGATGGTCCACGTC |
| circRNA | DNAJC3 | CCTGAAAGATCGAGCAGAGGCCTATTTGATAGAGGAAATGTATGATGAAGCTCAAATCTAATCCAAGTGAAAATGAAGAAAAGGAAGCACAGTCTCAACT |
| circRNA | ERBIN | ACTGGAGAGAACAAGTACTTCGACATATTGAAGCCAAAAAGTTAGAAAAGGCAAGAAGATGAAAATTTTAACAGCCTTTTACAAAATGGAGATATTTTAA |
| circRNA | ERC1 | AAATCACCATTTGGAAGGAACAGTACAGAGTTGTACAGGAGGAAAACCAGATATGGTGTAAGATACTTCTTCAAGATTGGACAGCTGGGGACCTTCTTCT |
| circRNA | EXOC6B | AGAGCATCCGCAAACATTCAGACAAAATTGGAGAGACTGCCATGAAGCAAAATCAAGTGACGGATACTAATAGAAAACTACAACATGAGGGAAAGGAACT |
| circRNA | FBXW7 | ACCTGCCCGTTCACCAACTCTCCTCCCCATTCTATACAAAAACAACAAAAGATTACTTCCTTAGGATAGATTGCCAGAAGTGGAGTTACTGGGTCAGAGG |
| circRNA | FNDC3B | AGAGCGACGAGCAAGAAGCAGCCCAAAGTCGAATGATTCAGACTTGCAAGGTGATTGAAGATAGTACTGGAGTCCGCCGGGTGGTGGTCACACCCCAGTC |
| mRNA | FUS | AGGTGGAGGTAACTATGGCCAAGATCAATCCTCCATGAGTAGTGGTGGTGGCAGTGGTGGCGGTTATGGCAATCAAGACCAGAGTGGTGGAGGTGGCAGC |
| mRNA | GAPDH | ACTGAATCTCCCCTCCTCACAGTTGCCATGTAGACCCCTTGAAGAGGGGAGGGGCCTAGGGAGCCGCACCTTGTCATGTACCATCAATAAAGTACCCTGT |
| circRNA | GDI2 | GTGGAAAAATCTACAAGGTTCCTTCCACTGAAGCAGAAGCCCTGGCATCTAGAATGTATCCTGTCAGGTATAATGTCAGTGAATGGCAAGAAAGTTCTTC |
| circRNA | GPATCH1 | CCTTAGTAAACAAAGAGGAAGAGCATGCACCAGAATTATCCGCAAATCAGTTCAACTTTAGTTGGCTTACCAAGAGTGAAGCGTGACAAGTACTCAGTCT |
| mRNA | GUSB | CCGATTTCATGACTGAACAGTCACCGACGAGAGTGCTGGGGAATAAAAAGGGGATCTTCACTCGGCAGAGACAACCAAAAAGTGCAGCGTTCCTTTTGCG |
| circRNA | HIPK3 | TGTATCAAAGACTGTTTGTTCAACATATCTACAATCTCGGTACTACAGGTATGGCCTCACAAGTCTTGGTCTACCCACCATATGTTTATCAAACTCAGTC |
| mRNA | HNRNPL | CCGAGTCTTCAATGTCTTCTGCTTATATGGCAATGTGGAGAAGGTGAAATTCATGAAAAGCAAGCCGGGGGCCGCCATGGTGGAGATGGCTGATGGCTAC |
| circRNA | KRT1 | CTTGTTAACCTTGGTGGCAGTAAAAGCATCTCCATAAGTGTGGCTAGAGGAGGTGGTGGGAGATTTTCAAGCTGTGGTGGTGGTGGTGGTAGCTTTGGTG |
| circRNA | MAN1A2_a | CTACTTGCAGCATATTACCTATCAGGAGAGGAGGGAAGAGGAAGAACGTCTGAGAAATAAAATTCGAGCTGATCATGAGAAGGCCTTGGAAGAAGCAAAA |
| circRNA | MITD1 | GATGGATGATTAAGATTGGAAGGGGACTTGATTATTTTAAGAAACCACAGCTGTATAACTTTCTTCGATTTTGTGAGATGCTTATTAAGAGACCATGTAA |
| circRNA | MORC3 | CGAAGAGTCTTGCCTACATCGAACGTGATGTTTATCGACCAAAATTTTTAATGATTAATTTAGCAGAATCAAAAGCCAGCCTTGCTGCAATTCTGGAACA |
| circRNA | PARD3 | TATGCCCAAGTCAAGAAGCCGCGGAATTCCAAACCCTCACCTGTAGACAGGTTTGGCAAACATCGAAAAGATGACAAGATTGAGAAAACGGGTAAAATAA |
| circRNA | PCMTD1 | CCTGGGAAGTGGAACCGGATATTTAAGTACAATGGTGGGCTTAATTTTAGGATTAATTTATTTTTGGAAATCAAGTGCAATATTGGAAGCCATTTCTACT |
| circRNA | PNN | GTGTCCAGCTACCCAAAAACTAATAGAAGAGTCACAGAGAAAAATGAACGCAAAAGCGGCGCCAGGAAATTGAACAAAAACTTGAAGTTCAGGCAGAAGA |
| circRNA | PRKD3 | GCCCAGGAACTGTCTTTATCTGCTGTCAAGGATCTTGTGTGCTCCATAGTTTATCAAAAGGCTAACTATATGTCAGAAAGCATCAGCTTTTCTCGATGGA |
| mRNA | QKI | CTTCTGCGGGATCTTCAACCACCTCGAGCGGCTGCTGGACGAAGAAATTAGCAGAGTACGGAAAGACATGTACAATGACACATTAAATGGCAGTACAGAG |
| circRNA | RBM23 | AACAGTGGCAATGAGACCAGTGGAAGCAGCACCATCGGGGAGACAAGCAAATCTGACAGGATGGCATCTGATGACTTTGACATAGTGATTGAGGCCATGC |
| circRNA | RBM33 | CTGATGAAGTGTTAGACATCGAGATCAATGAACCTTTAGATGAATTTACATGAACTTGAAGATGATTTACTTGGAGAAGATTTGCTATCTGGCAAAAAGA |
| mRNA | RPL19 | CCAATGCCCGAATGCCAGAGAAGGTCACATGGATGAGGAGAATGAGGATTTTGCGCCGGCTGCTCAGAAGATACCGTGAATCTAAGAAGATCGATCGCCA |
| mRNA | RPLP0 | CGAAATGTTTCATTGTGGGAGCAGACAATGTGGGCTCCAAGCAGATGCAGCAGATCCGCATGTCCCTTCGCGGGAAGGCTGTGGTGCTGATGGGCAAGAA |
| circRNA | SCMH1 | ACCCCAGGATGCTGCCACCATCCCCAGCTCAGCCATGCAGGCCCCAACAGTTGTGATTCCAAAGAATCCCTATCCTGCCTCCGATGTGAATACTGAGAAG |
| circRNA | SDF4 | GGTTGCCGACGCCATCAGGCTCAACGAGGAACTCAAAGTGGATGAGGAAAGGTGGATGTGAACACTGACCGGAAGATCAGTGCCAAGGAGATGCAGCGCT |
| circRNA | SLC8A1 | GGATTTTGAGGACACTTGTGGAGAGCTCGAATTCCAGAATGATGAAATTGTTAGGTTGTGACAGTTGGAAGTGTCATGTACAACATGCGGCGATTAAGTC |
| circRNA | SMAD2 | TGGAGAAACAAGTGACCAACAGTTGAATCAAAGTATGGACACAGGTTCGATACAAGAGGCTGTTTTCCTAGCGTGGCTTGCTGCCTTTGGTAAGAACATG |
| circRNA | SMARCA5 | AGTCAAGTGTTTATAACTTCGAAGGAGAAGACTATAGAGAAAAACAAAAGGGAGGCTTGTGGATCAGAATCTGAACAAAATTGGGAAAGATGAAATGCTT |
| circRNA | TRIM35 | CGGAGGTGCTGGCACATGAGATCGAGCGGCTGCAGATGGAGATGAAGGAGGACGACGTTTCTTTTCTCATGGTGGAGGCTGCATGGCTGGAAGGCCGGAT |
| circRNA | TULP4 | GAGTACTCCACAGAGGATAAATTTCAACCTCCGGGGCCACAATAGCGAGATTTGTAAGACTCCAGGGCCTCCCAGCCGTGAATAATCTGATGGTTCCTGA |
| circRNA | VCAN | AAGAGTCAGTGGAAGGCACGGCAATCTATTTACCAGGTCGAATGAGTGATTTGAGTGTAATTGGTCATCCAATAGATTCAGAATCTAAAGAAGATGAACC |
| circRNA | VPS16 | GGGGAATTCCAGCACCGCAGCAACCACTCCAGAAACTTTCCCTCCTTCAGATAAGCATGGATTCCTGGTTCATTCTTGTTCTGCTCGGCAGTGGTCTGAT |
| circRNA | YPEL2 | GCATTCACTGCAGAGCTCACTTGGCCAATCATGATGAACTAATTTCCAAGGCTGCTGAGAACTAGCCCTAGACCTCTGCGTGAGGGTTCTTCTGCCGAAG |
| circRNA | YY1AP1 | CCTGCAACCCCAATCTCAATCCGGAGGCCAGTAGCACCAGGATATGTCTTTTCCAAGATGAGATGGGATTCTCCAACATGGAAGATGATGGCCCAGAAGA |
| circRNA | ZC3H6 | CACTCTTGAGGGGCCAGCTGACCCACAGGCGGACGTTCCCAGGAGTTCTGAGAAGATGGCGAATTAGAAGATGGTGAAATAGACGATGCAGGATTTGAAG |
| circRNA | ZKSCAN1 | GTCCCACTTCAAACATTCGTCTCGGAAACCCCGCCTCTTACAGTCACGAGGAATAGTAAAGAAACACATCATAAAACCTCCCAGGACATAAAGGTGAGCA |
| circRNA | ZMYM4 | CACAGCAGGGACTACTAGACAAAATAAAAGATGAACCTGACAATGCTCAAGTGGTGGTATCATGGATACAGAAATGTCTGAAGATATAGACCACAACTTA |
| circRNA | ZNF609 | AGGAAGGGGAGAATGAGTGTCGCCTGCTAAAGAAAGTCAAGTCTGAAAAGCAATGATGTTGTCCACTGGGCATGTACTGACCAATGTGGCAGGTCTGAGA |
| circRNA | ZNF91 | AAGCATTACAAATATGAAGAGGCATTATTTATGACCTTTTCTATGGAAAGGTATATGTCCTCATTTTCCTCAAGACTTTTGGCCAGAGCAGAGCATGGAA |

**Supplementary Table 2.** The 128 high abundance circRNAs detected in the lesional skin biopsies listed according to average RPM.

| **Chromosome** | **start** | **end** | **sense gene** | **antisense gene** | **strand** | **average RPM** | **average CTL ratio** | **Found by**  **CircExplorer (0=no, 1= yes)** | **circBase ID** |
| --- | --- | --- | --- | --- | --- | --- | --- | --- | --- |
| chr1 | 1158623 | 1159348 | SDF4 | . | - | 1.608466 | 1.188625 | 1 | hsa_circ_0000002 |
| chr4 | 144464661 | 144465125 | SMARCA5 | . | + | 1.509605 | 0.980435 | 1 | hsa_circ_0001445 |
| chr4 | 36230203 | 36231267 | ARAP2 | . | - | 1.262622 | 0.572949 | 1 | hsa_circ_0069399 |
| chr1 | 117944807 | 117963271 | MAN1A2 | . | + | 1.135071 | 0.692023 | 1 | hsa_circ_0000118 |
| chr6 | 4891946 | 4892613 | CDYL | . | + | 1.129937 | 3.496164 | 1 | hsa_circ_0008285 |
| chr8 | 52773404 | 52773806 | PCMTD1 | . | - | 1.084443 | 0.962882 | 1 | hsa_circ_0001801 |
| chr15 | 64791491 | 64792365 | ZNF609 | . | + | 1.024965 | 13.03308 | 0 | hsa_circ_0000615 |
| chr19 | 33604672 | 33605325 | GPATCH1 | . | + | 0.887522 | 3.556228 | 1 | hsa_circ_0008287 |
| chr18 | 9182379 | 9221997 | ANKRD12 | . | + | 0.777933 | 2.706397 | 1 | hsa_circ_0000826 |
| chr21 | 37711076 | 37717005 | MORC3 | . | + | 0.722801 | 0.396759 | 0 | hsa_circ_0001189 |
| chrX | 139865339 | 139866824 | . | CDR1 | + | 0.693394 | 41 | 0 | hsa_circ_0001946 |
| chr14 | 23378691 | 23380612 | RBM23 | . | - | 0.595654 | 0.222039 | 1 | hsa_circ_0000524 |
| chr11 | 33307958 | 33309057 | HIPK3 | . | + | 0.506226 | 1.211131 | 1 | hsa_circ_0000284 |
| chr2 | 72945231 | 72960247 | EXOC6B | . | - | 0.483172 | 0.366678 | 1 | hsa_circ_0009043 |
| chr7 | 99621041 | 99621930 | ZKSCAN1 | . | + | 0.467026 | 1.048848 | 1 | hsa_circ_0001727 |
| chr4 | 87967317 | 87968746 | AFF1 | . | + | 0.457118 | 0.687955 | 1 | hsa_circ_0001423 |
| chr11 | 35222628 | 35226187 | CD44 | . | + | 0.445485 | 0.033452 | 1 | hsa_circ_0021727 |
| chr7 | 155465560 | 155473602 | RBM33 | . | + | 0.431601 | 0.817773 | 1 | hsa_circ_0001772 |
| chr1 | 117944807 | 117957453 | MAN1A2 | . | + | 0.394504 | 0.11411 | 1 | hsa_circ_0000117 |
| chr4 | 153332454 | 153333681 | FBXW7 | . | - | 0.385511 | 5.422619 | 1 | hsa_circ_0001451 |
| chr6 | 58774981 | 58778378 | . | . | - | 0.385495 | 28 | 0 | Not in circBase |
| chr1 | 155408117 | 155429689 | ASH1L | . | - | 0.376322 | 0.415177 | 1 | hsa_circ_0003247 |
| chr9 | 138773478 | 138774924 | CAMSAP1 | . | - | 0.366565 | 0.51142 | 1 | hsa_circ_0001900 |
| chr18 | 45391429 | 45423180 | SMAD2 | . | - | 0.35858 | 0.649195 | 1 | hsa_circ_0000847 |
| chr12 | 53073882 | 53074008 | KRT1 | . | - | 0.354944 | 2.522781 | 0 | Not in circBase |
| chr8 | 61653817 | 61655656 | CHD7 | . | + | 0.336956 | 1.302274 | 1 | hsa_circ_0084582 |
| chr1 | 117944807 | 117984947 | MAN1A2 | . | + | 0.334729 | 0.168827 | 1 | hsa_circ_0000119 |
| chr2 | 37543379 | 37544322 | PRKD3 | . | - | 0.317686 | 1.540873 | 0 | hsa_circ_0000992 |
| chr17 | 20107645 | 20109225 | SPECC1 | . | + | 0.317507 | 1.126797 | 1 | hsa_circ_0000745 |
| chr14 | 99924615 | 99932150 | SETD3 | . | - | 0.313469 | 0.7488 | 1 | hsa_circ_0000567 |
| chr12 | 1136913 | 1137738 | ERC1 | . | + | 0.306063 | 3.561651 | 1 | hsa_circ_0000373 |
| chr6 | 18236682 | 18258636 | DEK | . | - | 0.297317 | 0.07994 | 1 | hsa_circ_0075796 |
| chr1 | 35824525 | 35827390 | ZMYM4 | . | + | 0.296531 | 2.255927 | 1 | hsa_circ_0011536 |
| chr1 | 117944807 | 117948267 | MAN1A2 | . | + | 0.295046 | 0.100111 | 1 | hsa_circ_0000116 |
| chr13 | 96409897 | 96416207 | DNAJC3 | . | + | 0.290568 | 0.119477 | 1 | hsa_circ_0101041 |
| chr2 | 40655612 | 40657444 | SLC8A1 | . | - | 0.289039 | 0.805176 | 1 | hsa_circ_0000994 |
| chr5 | 122881110 | 122893258 | CSNK1G3 | . | + | 0.285558 | 0.605536 | 1 | hsa_circ_0001522 |
| chr1 | 41536266 | 41541123 | SCMH1 | . | - | 0.280304 | 2.898413 | 1 | hsa_circ_0000061 |
| chr10 | 5836847 | 5842668 | GDI2 | . | - | 0.272226 | 0.034101 | 1 | hsa_circ_0002665 |
| chr10 | 70719561 | 70720005 | DDX21 | . | + | 0.270347 | 0.055491 | 1 | hsa_circ_0008865 |
| chr17 | 57430575 | 57430887 | YPEL2 | . | + | 0.268954 | 4.040598 | 1 | hsa_circ_0005600 |
| chr14 | 39648294 | 39648666 | PNN | . | + | 0.26391 | 0.175994 | 1 | hsa_circ_0101802 |
| chr16 | 68155889 | 68160513 | NFATC3 | . | + | 0.26335 | 0.728524 | 1 | hsa_circ_0000711 |
| chr3 | 196118683 | 196129890 | UBXN7 | . | - | 0.252995 | 0.445781 | 1 | hsa_circ_0001380 |
| chr18 | 9204473 | 9221997 | ANKRD12 | . | + | 0.250402 | 0.515008 | 1 | hsa_circ_0003865 |
| chr10 | 128768965 | 128788867 | DOCK1 | . | + | 0.246693 | 0.114343 | 1 | hsa_circ_0007142 |
| chr9 | 33953282 | 33963789 | UBAP2 | . | - | 0.241049 | 0.496666 | 1 | hsa_circ_0001847 |
| chr19 | 47767859 | 47768203 | CCDC9 | . | + | 0.240812 | 2.459967 | 1 | hsa_circ_0000944 |
| chr6 | 158733082 | 158735300 | TULP4 | . | + | 0.231547 | 1.476768 | 0 | hsa_circ_0131202 |
| chr18 | 8718421 | 8720494 | MTCL1 | . | + | 0.228358 | 1.904243 | 1 | hsa_circ_0000825 |
| chr10 | 5815804 | 5842668 | GDI2 | . | - | 0.228114 | 0.032835 | 1 | hsa_circ_0017586 |
| chr12 | 27867712 | 27877119 | MRPS35 | . | + | 0.216372 | 0.205415 | 1 | hsa_circ_0000384 |
| chr4 | 48371865 | 48385801 | SLAIN2 | . | + | 0.21494 | 0.495204 | 1 | hsa_circ_0126525 |
| chr9 | 114148656 | 114154104 | KIAA0368 | . | - | 0.21129 | 0.115614 | 1 | hsa_circ_0001882 |
| chr10 | 34558584 | 34573173 | PARD3 | . | - | 0.211271 | 0.122517 | 1 | hsa_circ_0018168 |
| chr14 | 97299803 | 97327072 | VRK1 | . | + | 0.209406 | 0.270524 | 1 | hsa_circ_0000566 |
| chr1 | 200729966 | 200784772 | CAMSAP2 | . | + | 0.209184 | 0.250623 | 1 | hsa_circ_0015839 |
| chr3 | 56626997 | 56628056 | CCDC66 | . | + | 0.208025 | 0.615174 | 1 | hsa_circ_0001313 |
| chr5 | 65349233 | 65350779 | ERBIN | . | + | 0.205232 | 0.132339 | 1 | hsa_circ_0072732 |
| chr2 | 113057425 | 113089859 | ZC3H6 | . | + | 0.204389 | 1.640858 | 0 | hsa_circ_0117028 |
| chr3 | 71090478 | 71102924 | FOXP1 | . | - | 0.203217 | 0.31245 | 1 | hsa_circ_0008234 |
| chr11 | 130130750 | 130131824 | ZBTB44 | . | - | 0.200009 | 1.2024 | 1 | hsa_circ_0002484 |
| chr6 | 87925620 | 87928449 | ZNF292 | . | + | 0.19738 | 0.339853 | 1 | hsa_circ_0004058 |
| chr13 | 33091993 | 33101669 | N4BP2L2 | . | - | 0.185274 | 0.273379 | 1 | hsa_circ_0000471 |
| chr5 | 82832825 | 82838087 | VCAN | . | + | 0.184873 | 0.102642 | 1 | hsa_circ_0073237 |
| chr20 | 2944917 | 2945848 | VPS16 | . | + | 0.183745 | 0.259637 | 1 | hsa_circ_0006117 |
| chr18 | 19345732 | 19359646 | MIB1 | . | + | 0.182304 | 0.486833 | 1 | hsa_circ_0000835 |
| chr1 | 31465236 | 31468067 | PUM1 | . | - | 0.181372 | 0.05761 | 1 | hsa_circ_0000043 |
| chr9 | 16727794 | 16738483 | BNC2 | . | - | 0.178763 | 0.615467 | 1 | hsa_circ_0008732 |
| chr13 | 41515056 | 41518061 | ELF1 | . | - | 0.178716 | 0.076612 | 1 | hsa_circ_0030051 |
| chr9 | 33971648 | 33973235 | UBAP2 | . | - | 0.176861 | 0.936508 | 1 | hsa_circ_0001851 |
| chr4 | 73956383 | 73958017 | ANKRD17 | . | - | 0.175434 | 2.262626 | 1 | hsa_circ_0007883 |
| chr7 | 155457868 | 155473602 | RBM33 | . | + | 0.174215 | 0.376704 | 1 | hsa_circ_0001771 |
| chr20 | 62407030 | 62422143 | ZBTB46 | . | - | 0.172409 | 9.333333 | 1 | hsa_circ_0002805 |
| chr1 | 155644800 | 155649303 | YY1AP1 | . | - | 0.171883 | 0.478506 | 1 | hsa_circ_0003608 |
| chr20 | 46252654 | 46262380 | NCOA3 | . | + | 0.170871 | 0.17424 | 1 | hsa_circ_0001165 |
| chr16 | 3900297 | 3901010 | CREBBP | . | - | 0.170433 | 0.759133 | 1 | hsa_circ_0007637 |
| chr10 | 102683731 | 102685776 | SLF2 | . | + | 0.1691 | 0.25573 | 1 | hsa_circ_0006654 |
| chr18 | 44526019 | 44526886 | . | . | + | 0.16591 | 1.379941 | 1 | hsa_circ_0108513 |
| chr2 | 99786012 | 99787892 | MITD1 | . | - | 0.16512 | 0.711765 | 1 | hsa_circ_0001050 |
| chr12 | 70193988 | 70195501 | RAB3IP | . | + | 0.16416 | 0.933155 | 1 | hsa_circ_0000419 |
| chr17 | 57808781 | 57816308 | VMP1 | . | + | 0.163336 | 0.054722 | 1 | hsa_circ_0006508 |
| chr10 | 128768965 | 128798571 | DOCK1 | . | + | 0.161256 | 0.176187 | 1 | hsa_circ_0002669 |
| chr15 | 41961025 | 41962156 | MGA | . | + | 0.160549 | 0.37138 | 1 | hsa_circ_0000591 |
| chr11 | 107916996 | 107925682 | CUL5 | . | + | 0.160162 | 0.198592 | 1 | hsa_circ_0024169 |
| chr18 | 9195548 | 9221997 | ANKRD12 | . | + | 0.159151 | 0.108123 | 1 | hsa_circ_0046843 |
| chr4 | 178274461 | 178274882 | NEIL3 | . | + | 0.157907 | 0.457783 | 1 | hsa_circ_0001459 |
| chr10 | 28872327 | 28884970 | WAC | . | + | 0.1552 | 0.045228 | 1 | hsa_circ_0007503 |
| chr3 | 157839891 | 157841780 | RSRC1 | . | + | 0.154854 | 0.066848 | 1 | hsa_circ_0001355 |
| chr1 | 1586822 | 1650894 | CDK11B | MMP23A | - | 0.153833 | 1.199123 | 0 | hsa_circ_0000005 |
| chr1 | 85331067 | 85331821 | LPAR3 | . | - | 0.153784 | 4.016667 | 1 | hsa_circ_0004390 |
| chr8 | 130788347 | 130789837 | GSDMC | . | - | 0.153305 | 0.095012 | 1 | Not in circBase |
| chr21 | 17135209 | 17138460 | USP25 | . | + | 0.15089 | 0.144317 | 1 | hsa_circ_0001178 |
| chr3 | 31617887 | 31621588 | STT3B | . | + | 0.145883 | 0.185935 | 1 | hsa_circ_0001278 |
| chr15 | 93467550 | 93472321 | CHD2 | . | + | 0.144921 | 0.109581 | 1 | hsa_circ_0007262 |
| chr4 | 56277780 | 56284152 | TMEM165 | . | + | 0.142422 | 0.026855 | 1 | hsa_circ_0001414 |
| chr16 | 66764014 | 66766408 | DYNC1LI2 | . | - | 0.141032 | 0.16572 | 0 | hsa_circ_0000706 |
| chr3 | 171965322 | 171969331 | FNDC3B | . | + | 0.140287 | 0.417893 | 1 | hsa_circ_0006156 |
| chr4 | 37633006 | 37640126 | RELL1 | . | - | 0.140086 | 0.391597 | 1 | hsa_circ_0001400 |
| chr1 | 33760537 | 33760906 | ZNF362 | . | + | 0.135953 | 0.989226 | 1 | hsa_circ_0009027 |
| chr1 | 1747194 | 1770677 | GNB1 | . | - | 0.134765 | 0.168406 | 1 | hsa_circ_0008702 |
| chr12 | 95602618 | 95605043 | FGD6 | . | - | 0.133459 | 0.74663 | 1 | hsa_circ_0099549 |
| chr10 | 17746429 | 17747740 | STAM | . | + | 0.133237 | 0.091825 | 1 | hsa_circ_0008311 |
| chr4 | 2951660 | 2952972 | NOP14 | NOP14-AS1 | - | 0.132802 | 0.067988 | 1 | hsa_circ_0006737 |
| chr4 | 77065301 | 77065626 | NUP54 | . | - | 0.132599 | 0.136979 | 1 | hsa_circ_0070040 |
| chr10 | 70152894 | 70154208 | RUFY2 | . | - | 0.132079 | 0.157655 | 1 | hsa_circ_0000239 |
| chr22 | 22160138 | 22162135 | MAPK1 | . | - | 0.131724 | 0.11687 | 1 | hsa_circ_0008870 |
| chr2 | 168920009 | 168986268 | STK39 | . | - | 0.131128 | 0.873016 | 1 | hsa_circ_0005882 |
| chrX | 19983161 | 19988416 | CXorf23 | . | - | 0.130303 | 2.644444 | 0 | hsa_circ_0140068 |
| chr10 | 126631025 | 126631876 | ZRANB1 | . | + | 0.125109 | 0.583883 | 1 | hsa_circ_0000268 |
| chr2 | 242644067 | 242651486 | ING5 | . | + | 0.123946 | 0.153975 | 1 | hsa_circ_0001124 |
| chr11 | 120343758 | 120348235 | ARHGEF12 | . | + | 0.122029 | 0.020186 | 1 | hsa_circ_0009021 |
| chr16 | 80718434 | 80719026 | CDYL2 | . | - | 0.121158 | 0.401961 | 1 | hsa_circ_0004087 |
| chr4 | 187627716 | 187630999 | FAT1 | . | - | 0.120759 | 0.279702 | 1 | hsa_circ_0001461 |
| chr17 | 81042813 | 81043199 | METRNL | . | + | 0.1192 | 0.19228 | 1 | Not in circBase |
| chr10 | 128768965 | 128860040 | DOCK1 | . | + | 0.118466 | 0.193794 | 1 | hsa_circ_0020394 |
| chr14 | 31404368 | 31425448 | STRN3 | . | - | 0.118131 | 0.067419 | 1 | hsa_circ_0031446 |
| chr1 | 92798947 | 92846430 | RPAP2 | . | + | 0.117228 | 1.544444 | 1 | hsa_circ_0000091 |
| chr5 | 132227855 | 132228810 | AFF4 | . | - | 0.116723 | 0.07623 | 1 | hsa_circ_0001529 |
| chr16 | 18846213 | 18856973 | SMG1 | . | - | 0.112306 | 0.163112 | 1 | hsa_circ_0038258 |
| chr7 | 23650789 | 23651172 | CCDC126 | . | + | 0.109203 | 0.453373 | 1 | hsa_circ_0001684 |
| chr8 | 42761315 | 42798588 | HOOK3 | . | + | 0.106398 | 0.216678 | 1 | hsa_circ_0084143 |
| chr9 | 37126308 | 37126939 | ZCCHC7 | . | + | 0.105243 | 0.160809 | 1 | hsa_circ_0001860 |
| chr10 | 105197771 | 105198565 | PDCD11 | . | + | 0.104669 | 0.208155 | 1 | hsa_circ_0000258 |
| chr22 | 24698162 | 24709434 | SPECC1L | . | + | 0.10194 | 0.094009 | 1 | hsa_circ_0007312 |
| chr4 | 39915230 | 39927553 | PDS5A | . | - | 0.101171 | 0.047122 | 1 | hsa_circ_0007308 |
| chr6 | 76344422 | 76388643 | SENP6 | . | + | 0.101164 | 0.053543 | 1 | hsa_circ_0001613 |
| chr12 | 123983090 | 123984083 | RILPL1 | . | - | 0.093479 | 0.485294 | 1 | hsa_circ_0007552 |

**Supplementary Table 3.** The 285 high abundance circRNAs detected in the non-lesional skin biopsies listed according to average RPM.

| **Chromosome** | **start** | **end** | **sense gene** | **antisense gene** | **strand** | **average RPM** | **average CTL ratio** | **Found by**  **CircExplorer (0=no, 1= yes)** | **circBase ID** |
| --- | --- | --- | --- | --- | --- | --- | --- | --- | --- |
| chr4 | 144464661 | 144465125 | SMARCA5 | . | + | 3.746063 | 2.126652 | 1 | hsa_circ_0001445 |
| chr8 | 52773404 | 52773806 | PCMTD1 | . | - | 3.017865 | 1.154025 | 1 | hsa_circ_0001801 |
| chr1 | 117944807 | 117963271 | MAN1A2 | . | + | 2.556731 | 1.187509 | 1 | hsa_circ_0000118 |
| chr1 | 1158623 | 1159348 | SDF4 | . | - | 2.484468 | 1.352263 | 1 | hsa_circ_0000002 |
| chr15 | 64791491 | 64792365 | ZNF609 | . | + | 2.342102 | 28.60952 | 0 | hsa_circ_0000615 |
| chr21 | 37711076 | 37717005 | MORC3 | . | + | 2.146881 | 0.77296 | 0 | hsa_circ_0001189 |
| chr18 | 9182379 | 9221997 | ANKRD12 | . | + | 2.112748 | 3.813259 | 1 | hsa_circ_0000826 |
| chr19 | 33604672 | 33605325 | GPATCH1 | . | + | 2.008743 | 7.070943 | 1 | hsa_circ_0008287 |
| chr6 | 4891946 | 4892613 | CDYL | . | + | 1.994914 | 4.870296 | 1 | hsa_circ_0008285 |
| chr4 | 36230203 | 36231267 | ARAP2 | . | - | 1.90811 | 0.6022 | 1 | hsa_circ_0069399 |
| chrX | 139865339 | 139866824 | . | CDR1 | + | 1.801118 | 104.3333 | 0 | hsa_circ_0001946 |
| chr2 | 72945231 | 72960247 | EXOC6B | . | - | 1.733789 | 0.68526 | 1 | hsa_circ_0009043 |
| chr2 | 40655612 | 40657444 | SLC8A1 | . | - | 1.470716 | 1.441806 | 1 | hsa_circ_0000994 |
| chr14 | 23378691 | 23380612 | RBM23 | . | - | 1.065269 | 0.248806 | 1 | hsa_circ_0000524 |
| chr1 | 117944807 | 117957453 | MAN1A2 | . | + | 1.059937 | 0.191469 | 1 | hsa_circ_0000117 |
| chr11 | 35222628 | 35226187 | CD44 | . | + | 1.025952 | 0.073659 | 1 | hsa_circ_0021727 |
| chr7 | 155465560 | 155473602 | RBM33 | . | + | 1.001398 | 0.810639 | 1 | hsa_circ_0001772 |
| chr1 | 117944807 | 117984947 | MAN1A2 | . | + | 0.849133 | 0.423243 | 1 | hsa_circ_0000119 |
| chr17 | 57430575 | 57430887 | YPEL2 | . | + | 0.845706 | 5.587831 | 1 | hsa_circ_0005600 |
| chr1 | 35824525 | 35827390 | ZMYM4 | . | + | 0.824084 | 4.365734 | 1 | hsa_circ_0011536 |
| chr20 | 2944917 | 2945848 | VPS16 | . | + | 0.812817 | 0.828847 | 1 | hsa_circ_0006117 |
| chr8 | 27151596 | 27151827 | TRIM35 | . | - | 0.809614 | 1.532377 | 1 | hsa_circ_0083756 |
| chr11 | 33307958 | 33309057 | HIPK3 | . | + | 0.79605 | 1.499561 | 1 | hsa_circ_0000284 |
| chr9 | 138773478 | 138774924 | CAMSAP1 | . | - | 0.783238 | 0.782551 | 1 | hsa_circ_0001900 |
| chr2 | 113057425 | 113089859 | ZC3H6 | . | + | 0.770686 | 1.791489 | 0 | hsa_circ_0117028 |
| chr1 | 41536266 | 41541123 | SCMH1 | . | - | 0.768843 | 3.876587 | 1 | hsa_circ_0000061 |
| chr19 | 47767859 | 47768203 | CCDC9 | . | + | 0.753458 | 4.722854 | 1 | hsa_circ_0000944 |
| chr5 | 122881110 | 122893258 | CSNK1G3 | . | + | 0.746885 | 0.852149 | 1 | hsa_circ_0001522 |
| chr3 | 56626997 | 56628056 | CCDC66 | . | + | 0.743254 | 1.330314 | 1 | hsa_circ_0001313 |
| chr1 | 155644800 | 155649303 | YY1AP1 | . | - | 0.736515 | 1.167159 | 1 | hsa_circ_0003608 |
| chr1 | 117944807 | 117948267 | MAN1A2 | . | + | 0.722909 | 0.119527 | 1 | hsa_circ_0000116 |
| chr10 | 34558584 | 34573173 | PARD3 | . | - | 0.715073 | 0.213056 | 1 | hsa_circ_0018168 |
| chr4 | 87967317 | 87968746 | AFF1 | . | + | 0.699685 | 0.712061 | 1 | hsa_circ_0001423 |
| chr6 | 158733082 | 158735300 | TULP4 | . | + | 0.691981 | 1.169951 | 0 | hsa_circ_0131202 |
| chr13 | 96409897 | 96416207 | DNAJC3 | . | + | 0.688563 | 0.224521 | 1 | hsa_circ_0101041 |
| chr6 | 18236682 | 18258636 | DEK | . | - | 0.675798 | 0.127524 | 1 | hsa_circ_0075796 |
| chr14 | 39648294 | 39648666 | PNN | . | + | 0.668284 | 0.302878 | 1 | hsa_circ_0101802 |
| chr5 | 82832825 | 82838087 | VCAN | . | + | 0.663412 | 0.249595 | 1 | hsa_circ_0073237 |
| chr7 | 99621041 | 99621930 | ZKSCAN1 | . | + | 0.660294 | 0.88374 | 1 | hsa_circ_0001727 |
| chr2 | 99786012 | 99787892 | MITD1 | . | - | 0.657442 | 1.717135 | 1 | hsa_circ_0001050 |
| chr5 | 65349233 | 65350779 | ERBIN | . | + | 0.656291 | 0.224385 | 1 | hsa_circ_0072732 |
| chr3 | 171965322 | 171969331 | FNDC3B | . | + | 0.65011 | 0.97744 | 1 | hsa_circ_0006156 |
| chr10 | 70719561 | 70720005 | DDX21 | . | + | 0.644684 | 0.190164 | 1 | hsa_circ_0008865 |
| chr1 | 155408117 | 155429689 | ASH1L | . | - | 0.63584 | 0.427007 | 1 | hsa_circ_0003247 |
| chr11 | 107916996 | 107925682 | CUL5 | . | + | 0.635047 | 0.546534 | 1 | hsa_circ_0024169 |
| chr6 | 76344422 | 76388643 | SENP6 | . | + | 0.6215 | 0.23843 | 1 | hsa_circ_0001613 |
| chr8 | 618597 | 624047 | ERICH1 | . | - | 0.619075 | 1.031498 | 1 | hsa_circ_0136839 |
| chr9 | 37126308 | 37126939 | ZCCHC7 | . | + | 0.605946 | 0.447118 | 1 | hsa_circ_0001860 |
| chr4 | 153332454 | 153333681 | FBXW7 | . | - | 0.604605 | 8.064286 | 1 | hsa_circ_0001451 |
| chr2 | 168920009 | 168986268 | STK39 | . | - | 0.601119 | 1.652609 | 1 | hsa_circ_0005882 |
| chr15 | 41961025 | 41962156 | MGA | . | + | 0.601038 | 0.611925 | 1 | hsa_circ_0000591 |
| chr8 | 37971709 | 37976881 | ASH2L | . | + | 0.590923 | 1.550286 | 1 | hsa_circ_0001790 |
| chr12 | 1136913 | 1137738 | ERC1 | . | + | 0.59045 | 1.531823 | 1 | hsa_circ_0000373 |
| chr2 | 37543379 | 37544322 | PRKD3 | . | - | 0.577151 | 1.501635 | 0 | hsa_circ_0000992 |
| chr10 | 5836847 | 5842668 | GDI2 | . | - | 0.575988 | 0.052514 | 1 | hsa_circ_0002665 |
| chr3 | 157839891 | 157841780 | RSRC1 | . | + | 0.563835 | 0.167942 | 1 | hsa_circ_0001355 |
| chr4 | 48371865 | 48385801 | SLAIN2 | . | + | 0.535349 | 0.729214 | 1 | hsa_circ_0126525 |
| chr7 | 155457868 | 155473602 | RBM33 | . | + | 0.522924 | 0.551496 | 1 | hsa_circ_0001771 |
| chr6 | 87925620 | 87928449 | ZNF292 | . | + | 0.513707 | 0.496375 | 1 | hsa_circ_0004058 |
| chr14 | 99924615 | 99932150 | SETD3 | . | - | 0.507134 | 1.391732 | 1 | hsa_circ_0000567 |
| chr13 | 41515056 | 41518061 | ELF1 | . | - | 0.480026 | 0.124977 | 1 | hsa_circ_0030051 |
| chr18 | 9204473 | 9221997 | ANKRD12 | . | + | 0.476837 | 0.299959 | 1 | hsa_circ_0003865 |
| chr18 | 45391429 | 45423180 | SMAD2 | . | - | 0.471161 | 0.459374 | 1 | hsa_circ_0000847 |
| chr3 | 196118683 | 196129890 | UBXN7 | . | - | 0.469293 | 0.541541 | 1 | hsa_circ_0001380 |
| chr10 | 105767934 | 105778666 | SLK | . | + | 0.469114 | 0.047388 | 1 | hsa_circ_0000259 |
| chr5 | 167915606 | 167921655 | RARS | . | + | 0.454761 | 0.085171 | 1 | hsa_circ_0001550 |
| chr2 | 24357988 | 24369956 | FAM228B | . | + | 0.446377 | 4.669444 | 1 | hsa_circ_0000982 |
| chr10 | 128768965 | 128788867 | DOCK1 | . | + | 0.438201 | 0.113105 | 1 | hsa_circ_0007142 |
| chr1 | 33760537 | 33760906 | ZNF362 | . | + | 0.434854 | 1.938095 | 1 | hsa_circ_0009027 |
| chr14 | 31404368 | 31425448 | STRN3 | . | - | 0.431937 | 0.275788 | 1 | hsa_circ_0031446 |
| chr9 | 33953282 | 33963789 | UBAP2 | . | - | 0.429453 | 0.559118 | 1 | hsa_circ_0001847 |
| chr12 | 123983090 | 123984083 | RILPL1 | . | - | 0.427432 | 2.080303 | 1 | hsa_circ_0007552 |
| chr17 | 20107645 | 20109225 | SPECC1 | . | + | 0.424977 | 1.84954 | 1 | hsa_circ_0000745 |
| chr14 | 23375403 | 23380612 | RBM23 | . | - | 0.423633 | 0.131477 | 1 | hsa_circ_0031241 |
| chr7 | 99090662 | 99092254 | ZNF394 | . | - | 0.422579 | 2.897661 | 0 | hsa_circ_0001726 |
| chr4 | 37633006 | 37640126 | RELL1 | . | - | 0.415588 | 0.359851 | 1 | hsa_circ_0001400 |
| chr5 | 132227855 | 132228810 | AFF4 | . | - | 0.414911 | 0.186111 | 1 | hsa_circ_0001529 |
| chr4 | 148778703 | 148803083 | ARHGAP10 | . | + | 0.414261 | 0.474387 | 1 | hsa_circ_0071106 |
| chr10 | 128768965 | 128798571 | DOCK1 | . | + | 0.404371 | 0.142714 | 1 | hsa_circ_0002669 |
| chr8 | 42761315 | 42798588 | HOOK3 | . | + | 0.404192 | 0.475448 | 1 | hsa_circ_0084143 |
| chr16 | 18852886 | 18856973 | SMG1 | . | - | 0.402943 | 0.159832 | 1 | hsa_circ_0006434 |
| chr18 | 9195548 | 9221997 | ANKRD12 | . | + | 0.3993 | 0.130929 | 1 | hsa_circ_0046843 |
| chr16 | 68155889 | 68160513 | NFATC3 | . | + | 0.39763 | 0.691792 | 1 | hsa_circ_0000711 |
| chr16 | 53288349 | 53308214 | CHD9 | . | + | 0.394921 | 0.623994 | 1 | hsa_circ_0000702 |
| chr12 | 122825299 | 122826244 | CLIP1 | . | - | 0.394048 | 0.101824 | 1 | hsa_circ_0029069 |
| chr10 | 91511102 | 91522592 | KIF20B | . | + | 0.393814 | 0.684997 | 1 | hsa_circ_0019079 |
| chr17 | 65941524 | 65972074 | BPTF | . | + | 0.392244 | 0.870503 | 1 | hsa_circ_0000799 |
| chr15 | 62299506 | 62306191 | VPS13C | . | - | 0.386291 | 0.280764 | 1 | hsa_circ_0000607 |
| chr15 | 41036244 | 41037457 | RMDN3 | . | - | 0.385542 | 1.515986 | 1 | hsa_circ_0004942 |
| chr21 | 30693541 | 30702014 | BACH1 | . | + | 0.384998 | 0.259566 | 1 | hsa_circ_0001181 |
| chr13 | 33091993 | 33101669 | N4BP2L2 | . | - | 0.383116 | 0.262416 | 1 | hsa_circ_0000471 |
| chr15 | 52161413 | 52194233 | TMOD3 | . | + | 0.379449 | 0.102656 | 1 | hsa_circ_0035292 |
| chr22 | 22160138 | 22162135 | MAPK1 | . | - | 0.376306 | 0.212481 | 1 | hsa_circ_0008870 |
| chr10 | 17746429 | 17747740 | STAM | . | + | 0.372303 | 0.211126 | 1 | hsa_circ_0008311 |
| chr9 | 33971648 | 33973235 | UBAP2 | . | - | 0.371215 | 1.04462 | 1 | hsa_circ_0001851 |
| chr10 | 102683731 | 102685776 | SLF2 | . | + | 0.371068 | 0.267311 | 1 | hsa_circ_0006654 |
| chr4 | 129913321 | 129925031 | SCLT1 | . | - | 0.370821 | 1.129753 | 1 | hsa_circ_0001439 |
| chr12 | 46319924 | 46322642 | SCAF11 | . | - | 0.36931 | 0.275577 | 1 | hsa_circ_0025967 |
| chr20 | 62407030 | 62422143 | ZBTB46 | . | - | 0.366238 | 18.38889 | 1 | hsa_circ_0002805 |
| chr18 | 19345732 | 19359646 | MIB1 | . | + | 0.366173 | 0.353463 | 1 | hsa_circ_0000835 |
| chr7 | 30590251 | 30614497 | LOC401320 | . | - | 0.364231 | 1.579675 | 0 | hsa_circ_0134091 |
| chr8 | 48308935 | 48320523 | SPIDR | . | + | 0.360653 | 1.073076 | 1 | hsa_circ_0001798 |
| chr11 | 92085261 | 92088570 | FAT3 | . | + | 0.359283 | 6.305556 | 1 | hsa_circ_0000348 |
| chr8 | 61653817 | 61655656 | CHD7 | . | + | 0.358949 | 0.808367 | 1 | hsa_circ_0084582 |
| chr11 | 130130750 | 130131824 | ZBTB44 | . | - | 0.354223 | 1.287388 | 1 | hsa_circ_0002484 |
| chr7 | 23650789 | 23651172 | CCDC126 | . | + | 0.353612 | 0.828899 | 1 | hsa_circ_0001684 |
| chr20 | 34317233 | 34320057 | RBM39 | . | - | 0.349189 | 0.06741 | 1 | hsa_circ_0008817 |
| chr10 | 5741487 | 5756170 | FAM208B | . | + | 0.348124 | 1.512358 | 0 | hsa_circ_0000209 |
| chr19 | 8520288 | 8528570 | HNRNPM | . | + | 0.345525 | 0.14377 | 1 | hsa_circ_0006382 |
| chr12 | 70193988 | 70195501 | RAB3IP | . | + | 0.344843 | 1.355994 | 1 | hsa_circ_0000419 |
| chr11 | 68318588 | 68331900 | PPP6R3 | . | + | 0.339652 | 0.164259 | 1 | hsa_circ_0007660 |
| chr10 | 7839009 | 7844817 | ATP5C1 | . | + | 0.337539 | 0.688131 | 1 | hsa_circ_0007292 |
| chr3 | 138289159 | 138291774 | CEP70 | . | - | 0.333778 | 0.905839 | 1 | hsa_circ_0004524 |
| chr15 | 41988272 | 41991357 | MGA | . | + | 0.333559 | 0.299654 | 1 | hsa_circ_0000592 |
| chr17 | 45479497 | 45492285 | EFCAB13 | . | + | 0.328281 | 1.943878 | 1 | hsa_circ_0000778 |
| chr5 | 80071512 | 80088663 | MSH3 | . | + | 0.328247 | 0.199052 | 1 | hsa_circ_0001508 |
| chr3 | 169863210 | 169867032 | PHC3 | . | - | 0.328154 | 0.348956 | 1 | hsa_circ_0001360 |
| chr3 | 125032151 | 125050082 | ZNF148 | . | - | 0.32026 | 0.082668 | 1 | hsa_circ_0001333 |
| chr8 | 142264087 | 142264728 | SLC45A4 | LOC105375787 | - | 0.316396 | 10.77778 | 0 | hsa_circ_0001829 |
| chr2 | 68717321 | 68772444 | APLF | . | + | 0.316278 | 1.421741 | 1 | hsa_circ_0001023 |
| chr16 | 66764014 | 66766408 | DYNC1LI2 | . | - | 0.31499 | 0.207808 | 0 | hsa_circ_0000706 |
| chr5 | 137320945 | 137324004 | FAM13B | . | - | 0.314198 | 0.340889 | 1 | hsa_circ_0001535 |
| chr1 | 200729966 | 200784772 | CAMSAP2 | . | + | 0.314005 | 0.30706 | 1 | hsa_circ_0015839 |
| chr4 | 87685745 | 87689129 | PTPN13 | . | + | 0.312633 | 0.10439 | 1 | hsa_circ_0007948 |
| chr2 | 200233327 | 200298237 | SATB2 | . | - | 0.307914 | 3.514286 | 1 | hsa_circ_0003915 |
| chr21 | 17135209 | 17138460 | USP25 | . | + | 0.307674 | 0.156308 | 1 | hsa_circ_0001178 |
| chr2 | 113057425 | 113082774 | ZC3H6 | . | + | 0.3069 | 0.249372 | 1 | hsa_circ_0117027 |
| chrY | 2821949 | 2829687 | ZFY | . | + | 0.306556 | 0.864943 | 1 | hsa_circ_0001953 |
| chr5 | 63621012 | 63626233 | RNF180 | . | + | 0.304771 | 2.048148 | 1 | hsa_circ_0129323 |
| chr1 | 52981562 | 52992045 | ZCCHC11 | . | - | 0.302859 | 0.184639 | 1 | hsa_circ_0012553 |
| chr2 | 160194077 | 160252345 | BAZ2B | . | - | 0.302122 | 0.532868 | 1 | hsa_circ_0056829 |
| chr14 | 64465631 | 64489581 | SYNE2 | MIR548AZ | + | 0.301416 | 0.058772 | 1 | hsa_circ_0102377 |
| chr1 | 1586822 | 1650894 | CDK11B | MMP23A | - | 0.299941 | 1.902422 | 0 | hsa_circ_0000005 |
| chr15 | 93467550 | 93472321 | CHD2 | . | + | 0.298275 | 0.089042 | 1 | hsa_circ_0007262 |
| chr9 | 98740342 | 98766983 | ERCC6L2 | . | + | 0.297704 | 0.266693 | 1 | hsa_circ_0008720 |
| chr1 | 21295966 | 21329301 | EIF4G3 | . | - | 0.2965 | 0.142501 | 1 | hsa_circ_0005604 |
| chr3 | 43341245 | 43345284 | SNRK | . | + | 0.296281 | 0.688952 | 1 | hsa_circ_0004089 |
| chr4 | 39739039 | 39776553 | UBE2K | . | + | 0.296175 | 0.097462 | 1 | hsa_circ_0002590 |
| chr9 | 114148656 | 114154104 | KIAA0368 | . | - | 0.291796 | 0.115036 | 1 | hsa_circ_0001882 |
| chr10 | 28872327 | 28884970 | WAC | . | + | 0.288643 | 0.060462 | 1 | hsa_circ_0007503 |
| chr12 | 109046047 | 109048186 | CORO1C | . | - | 0.28702 | 0.442815 | 1 | hsa_circ_0000437 |
| chr12 | 27867712 | 27877119 | MRPS35 | . | + | 0.286405 | 0.203913 | 1 | hsa_circ_0000384 |
| chr4 | 48371865 | 48396670 | SLAIN2 | . | + | 0.286161 | 0.598087 | 0 | hsa_circ_0126526 |
| chr9 | 127670655 | 127674305 | GOLGA1 | . | - | 0.28013 | 0.172445 | 1 | hsa_circ_0002883 |
| chr1 | 67356836 | 67371058 | WDR78 | . | - | 0.279676 | 1.896825 | 1 | hsa_circ_0006677 |
| chr9 | 128419929 | 128434922 | MAPKAP1 | . | - | 0.279178 | 0.120053 | 1 | hsa_circ_0001890 |
| chr1 | 21083658 | 21100103 | HP1BP3 | . | - | 0.27715 | 0.135297 | 1 | hsa_circ_0000024 |
| chr11 | 120343758 | 120348235 | ARHGEF12 | . | + | 0.274065 | 0.029105 | 1 | hsa_circ_0009021 |
| chr4 | 129857809 | 129880932 | SCLT1 | . | - | 0.273979 | 0.276385 | 1 | hsa_circ_0070959 |
| chr15 | 63988322 | 64008672 | HERC1 | . | - | 0.273671 | 0.362309 | 1 | hsa_circ_0035796 |
| chr2 | 61343113 | 61345251 | KIAA1841 | . | + | 0.269814 | 0.701317 | 1 | hsa_circ_0007793 |
| chr10 | 95140975 | 95148911 | MYOF | . | - | 0.268933 | 0.14825 | 1 | hsa_circ_0005898 |
| chr19 | 13039155 | 13039661 | FARSA | . | - | 0.268382 | 1.856061 | 1 | hsa_circ_0000896 |
| chrX | 19983161 | 19988416 | CXorf23 | . | - | 0.267321 | 2.832906 | 0 | hsa_circ_0140068 |
| chr2 | 231307651 | 231314970 | SP100 | . | + | 0.266161 | 0.192518 | 1 | hsa_circ_0003922 |
| chr1 | 243579003 | 243589860 | SDCCAG8 | . | + | 0.265097 | 0.421605 | 1 | hsa_circ_0017241 |
| chr13 | 26974589 | 26975761 | CDK8 | . | + | 0.264802 | 0.768969 | 1 | hsa_circ_0003489 |
| chr16 | 80718434 | 80719026 | CDYL2 | . | - | 0.264315 | 0.845636 | 1 | hsa_circ_0004087 |
| chr17 | 57808781 | 57816308 | VMP1 | . | + | 0.263909 | 0.076334 | 1 | hsa_circ_0006508 |
| chr11 | 35222628 | 35229753 | CD44 | . | + | 0.263726 | 0.024277 | 1 | hsa_circ_0021728 |
| chr19 | 24014398 | 24016323 | . | . | + | 0.262314 | 1.296825 | 0 | hsa_circ_0109327 |
| chr3 | 160131260 | 160132305 | SMC4 | . | + | 0.262096 | 0.133774 | 1 | hsa_circ_0001356 |
| chr20 | 46252654 | 46262380 | NCOA3 | . | + | 0.253376 | 0.182672 | 1 | hsa_circ_0001165 |
| chr6 | 161455290 | 161471011 | MAP3K4 | . | + | 0.252719 | 0.860978 | 1 | hsa_circ_0078617 |
| chr1 | 58971731 | 59004982 | OMA1 | . | - | 0.252597 | 0.644776 | 0 | hsa_circ_0002316 |
| chr4 | 187627716 | 187630999 | FAT1 | . | - | 0.251826 | 0.170094 | 1 | hsa_circ_0001461 |
| chr3 | 169854206 | 169867032 | PHC3 | . | - | 0.250156 | 0.396112 | 1 | hsa_circ_0001359 |
| chr12 | 124904502 | 124915333 | NCOR2 | . | - | 0.248001 | 0.524733 | 1 | hsa_circ_0029308 |
| chr8 | 109462051 | 109462721 | EMC2 | . | + | 0.247736 | 0.046998 | 1 | hsa_circ_0135458 |
| chr6 | 71185112 | 71212494 | FAM135A | . | + | 0.246495 | 0.203461 | 1 | hsa_circ_0076961 |
| chr15 | 59204761 | 59209198 | SLTM | . | - | 0.246489 | 0.392063 | 1 | hsa_circ_0000605 |
| chr13 | 24164288 | 24200931 | TNFRSF19 | . | + | 0.245398 | 0.164244 | 1 | hsa_circ_0008784 |
| chr3 | 71090478 | 71102924 | FOXP1 | . | - | 0.245144 | 0.286287 | 1 | hsa_circ_0008234 |
| chrX | 53430497 | 53430825 | SMC1A | . | - | 0.244753 | 0.132675 | 1 | hsa_circ_0001921 |
| chr22 | 32874967 | 32875262 | FBXO7 | . | + | 0.24468 | 0.243085 | 1 | hsa_circ_0008832 |
| chr10 | 70152894 | 70154208 | RUFY2 | . | - | 0.243937 | 0.164545 | 1 | hsa_circ_0000239 |
| chr2 | 239090705 | 239093928 | ILKAP | . | - | 0.242719 | 1.759444 | 1 | hsa_circ_0001116 |
| chr12 | 112371689 | 112381173 | TMEM116 | . | - | 0.240094 | 0.180758 | 1 | hsa_circ_0004206 |
| chr7 | 104714065 | 104717880 | KMT2E | . | + | 0.239999 | 0.137736 | 1 | hsa_circ_0001736 |
| chr10 | 128768965 | 128860040 | DOCK1 | . | + | 0.239877 | 0.225478 | 1 | hsa_circ_0020394 |
| chr18 | 44526019 | 44526886 | . | . | + | 0.238816 | 4.560185 | 1 | hsa_circ_0108513 |
| chr1 | 231930987 | 231954263 | TSNAX-DISC1 | DISC2 | + | 0.237754 | 2.857937 | 1 | hsa_circ_0007848 |
| chr7 | 17908029 | 17915413 | SNX13 | . | - | 0.23712 | 0.097883 | 1 | hsa_circ_0005519 |
| chr9 | 138741982 | 138758382 | CAMSAP1 | . | - | 0.236835 | 0.279781 | 1 | hsa_circ_0089489 |
| chr14 | 67736417 | 67770316 | MPP5 | . | + | 0.236834 | 0.706445 | 1 | hsa_circ_0032261 |
| chr4 | 73956383 | 73958017 | ANKRD17 | . | - | 0.23633 | 0.992059 | 1 | hsa_circ_0007883 |
| chr2 | 203817281 | 203820481 | CARF | . | + | 0.235787 | 0.468028 | 1 | hsa_circ_0004919 |
| chr9 | 5968018 | 5988545 | KIAA2026 | . | - | 0.234871 | 0.234991 | 1 | hsa_circ_0138872 |
| chr1 | 174241551 | 174274265 | RABGAP1L | . | + | 0.233719 | 0.207935 | 1 | hsa_circ_0005089 |
| chr9 | 19286766 | 19305525 | DENND4C | . | + | 0.233347 | 0.117894 | 0 | hsa_circ_0005684 |
| chr7 | 91924202 | 91957214 | ANKIB1 | . | + | 0.232941 | 0.276013 | 1 | hsa_circ_0135062 |
| chr6 | 117010482 | 117026323 | KPNA5 | . | + | 0.232342 | 0.324039 | 1 | hsa_circ_0130438 |
| chr14 | 97299803 | 97327072 | VRK1 | . | + | 0.231989 | 0.245428 | 1 | hsa_circ_0000566 |
| chr1 | 35846859 | 35855699 | ZMYM4 | . | + | 0.230954 | 0.514409 | 1 | hsa_circ_0011542 |
| chr18 | 8718421 | 8720494 | MTCL1 | . | + | 0.230251 | 1.110804 | 1 | hsa_circ_0000825 |
| chr8 | 18622958 | 18662408 | PSD3 | . | - | 0.229815 | 0.195427 | 1 | hsa_circ_0002111 |
| chr6 | 77981056 | 77983024 | . | . | - | 0.229171 | 2.550311 | 0 | hsa_circ_0132345 |
| chr18 | 54423813 | 54426184 | WDR7 | . | + | 0.228736 | 0.399962 | 1 | hsa_circ_0000852 |
| chr6 | 13639794 | 13644961 | RANBP9 | . | - | 0.228314 | 0.029095 | 1 | hsa_circ_0001578 |
| chr8 | 130788347 | 130789837 | GSDMC | . | - | 0.227577 | 0.187593 | 1 | Not in circBase |
| chr6 | 76331247 | 76344527 | SENP6 | . | + | 0.226897 | 0.261371 | 1 | hsa_circ_0077078 |
| chr7 | 65705311 | 65751696 | TPST1 | . | + | 0.22535 | 0.475472 | 1 | hsa_circ_0006041 |
| chr21 | 17205666 | 17214859 | USP25 | . | + | 0.224137 | 0.110922 | 1 | hsa_circ_0005238 |
| chr2 | 136432901 | 136437894 | R3HDM1 | . | + | 0.223478 | 0.238761 | 1 | hsa_circ_0001070 |
| chr8 | 141874410 | 141900868 | PTK2 | . | - | 0.222874 | 0.243727 | 1 | hsa_circ_0002483 |
| chr16 | 11114049 | 11154879 | CLEC16A | . | + | 0.220878 | 1.092664 | 1 | hsa_circ_0000672 |
| chrX | 44383247 | 44386611 | FUNDC1 | . | - | 0.218406 | 0.229365 | 0 | hsa_circ_0007290 |
| chr1 | 8601272 | 8617582 | RERE | . | - | 0.218335 | 1.072569 | 1 | hsa_circ_0002158 |
| chr3 | 171969049 | 172028671 | FNDC3B | . | + | 0.217853 | 0.098647 | 1 | hsa_circ_0003692 |
| chr6 | 57058639 | 57075243 | RAB23 | . | - | 0.217694 | 0.21199 | 1 | hsa_circ_0004119 |
| chr1 | 108690900 | 108703915 | SLC25A24 | . | - | 0.217049 | 0.83076 | 1 | hsa_circ_0004270 |
| chr14 | 31602443 | 31602881 | HECTD1 | . | - | 0.216786 | 0.027936 | 1 | hsa_circ_0031485 |
| chrX | 84558411 | 84563222 | POF1B | . | - | 0.215641 | 0.00455 | 1 | hsa_circ_0091187 |
| chr11 | 34978930 | 34999729 | PDHX | . | + | 0.213055 | 0.139672 | 1 | hsa_circ_0021712 |
| chr15 | 44624185 | 44630515 | CASC4 | . | + | 0.212825 | 0.136905 | 1 | hsa_circ_0000596 |
| chr12 | 46622935 | 46637097 | SLC38A1 | . | - | 0.211108 | 0.098131 | 1 | hsa_circ_0000396 |
| chrX | 84600865 | 84622771 | POF1B | . | - | 0.210952 | 0.006646 | 1 | Not in circBase |
| chr1 | 31465236 | 31468067 | PUM1 | . | - | 0.209383 | 0.065946 | 1 | hsa_circ_0000043 |
| chr8 | 68044185 | 68049838 | CSPP1 | . | + | 0.20901 | 0.335953 | 1 | hsa_circ_0003388 |
| chr8 | 141856358 | 141874498 | PTK2 | . | - | 0.208549 | 0.172755 | 1 | hsa_circ_0002162 |
| chr2 | 211018218 | 211019335 | KANSL1L | . | - | 0.207901 | 0.588384 | 1 | hsa_circ_0008459 |
| chr1 | 85331067 | 85331821 | LPAR3 | . | - | 0.207402 | 4.329365 | 1 | hsa_circ_0004390 |
| chr10 | 112723882 | 112745523 | SHOC2 | . | + | 0.207081 | 0.111439 | 1 | hsa_circ_0020028 |
| chr19 | 38631823 | 38633350 | SIPA1L3 | . | + | 0.207045 | 1.375325 | 1 | hsa_circ_0006670 |
| chr7 | 27824781 | 27839709 | TAX1BP1 | . | + | 0.206179 | 0.016598 | 1 | hsa_circ_0134036 |
| chr7 | 158672374 | 158684024 | WDR60 | . | + | 0.204987 | 0.284513 | 1 | hsa_circ_0083226 |
| chr20 | 57014000 | 57016139 | VAPB | . | + | 0.203719 | 0.151782 | 1 | hsa_circ_0001173 |
| chr3 | 107429298 | 107435696 | BBX | . | + | 0.203612 | 0.193049 | 1 | hsa_circ_0001324 |
| chr4 | 129857809 | 129891623 | SCLT1 | . | - | 0.203039 | 0.528662 | 1 | hsa_circ_0070960 |
| chr11 | 118422500 | 118430579 | IFT46 | . | - | 0.202524 | 0.59843 | 1 | hsa_circ_0007372 |
| chr14 | 31416295 | 31425448 | STRN3 | . | - | 0.20195 | 0.074466 | 1 | hsa_circ_0031447 |
| chr10 | 105197771 | 105198565 | PDCD11 | . | + | 0.201831 | 0.204082 | 1 | hsa_circ_0000258 |
| chr14 | 102661274 | 102676199 | WDR20 | . | + | 0.200758 | 0.971164 | 1 | hsa_circ_0002553 |
| chr1 | 67423741 | 67428843 | MIER1 | . | + | 0.200227 | 0.078673 | 1 | hsa_circ_0113954 |
| chr4 | 56277780 | 56284152 | TMEM165 | . | + | 0.199832 | 0.054866 | 1 | hsa_circ_0001414 |
| chr4 | 178274461 | 178274882 | NEIL3 | . | + | 0.199106 | 0.804181 | 1 | hsa_circ_0001459 |
| chr11 | 85733409 | 85742653 | PICALM | . | - | 0.198809 | 0.04968 | 1 | hsa_circ_0023942 |
| chr10 | 15875628 | 15889942 | FAM188A | . | - | 0.198323 | 0.150361 | 1 | hsa_circ_0006665 |
| chr9 | 33948371 | 33956144 | UBAP2 | . | - | 0.197337 | 0.510826 | 1 | hsa_circ_0007367 |
| chr1 | 92798947 | 92846430 | RPAP2 | . | + | 0.197156 | 1.100649 | 1 | hsa_circ_0000091 |
| chr8 | 18656804 | 18662408 | PSD3 | . | - | 0.196937 | 0.195581 | 1 | hsa_circ_0004458 |
| chrX | 10031484 | 10066619 | WWC3 | . | + | 0.196585 | 1.37672 | 1 | hsa_circ_0001910 |
| chr13 | 51501542 | 51523641 | RNASEH2B | . | + | 0.196386 | 0.1496 | 1 | hsa_circ_0000489 |
| chr12 | 116668337 | 116675510 | MED13L | . | - | 0.195547 | 0.433958 | 0 | hsa_circ_0000443 |
| chr11 | 35222628 | 35231601 | CD44 | . | + | 0.194291 | 0.016199 | 1 | hsa_circ_0021729 |
| chr11 | 120345268 | 120348235 | ARHGEF12 | . | + | 0.194099 | 0.029959 | 1 | hsa_circ_0002100 |
| chr4 | 39915230 | 39927553 | PDS5A | . | - | 0.193052 | 0.071306 | 1 | hsa_circ_0007308 |
| chr3 | 3178943 | 3186394 | TRNT1 | . | + | 0.192565 | 0.712001 | 1 | hsa_circ_0123486 |
| chr10 | 126631025 | 126631876 | ZRANB1 | . | + | 0.191047 | 0.297774 | 1 | hsa_circ_0000268 |
| chr9 | 78682870 | 78722267 | PCSK5 | . | + | 0.190525 | 0.235927 | 1 | hsa_circ_0087234 |
| chr2 | 168869143 | 168873605 | STK39 | . | - | 0.189652 | 1.358153 | 1 | hsa_circ_0117963 |
| chr13 | 77779390 | 77818086 | MYCBP2 | . | - | 0.188092 | 0.03007 | 1 | hsa_circ_0030509 |
| chr9 | 33986757 | 34017187 | UBAP2 | . | - | 0.188023 | 0.359865 | 1 | hsa_circ_0086735 |
| chr4 | 73950965 | 73958017 | ANKRD17 | . | - | 0.187931 | 0.640383 | 1 | hsa_circ_0001417 |
| chr22 | 22153300 | 22162135 | MAPK1 | . | - | 0.187507 | 0.156727 | 1 | hsa_circ_0004872 |
| chr2 | 99802639 | 99812219 | MRPL30 | . | + | 0.18595 | 0.092966 | 1 | hsa_circ_0001051 |
| chr14 | 31398406 | 31425448 | STRN3 | . | - | 0.185581 | 0.15719 | 1 | hsa_circ_0101574 |
| chr7 | 22999874 | 23030758 | FAM126A | . | - | 0.185346 | 0.335161 | 1 | hsa_circ_0008951 |
| chr8 | 68163533 | 68172161 | ARFGEF1 | . | - | 0.184454 | 0.132043 | 1 | hsa_circ_0001807 |
| chr15 | 57743697 | 57754090 | CGNL1 | . | + | 0.183469 | 0.248214 | 1 | hsa_circ_0035435 |
| chr9 | 79996891 | 80022523 | VPS13A | . | + | 0.182342 | 0.056028 | 1 | hsa_circ_0008075 |
| chr11 | 74500670 | 74528759 | RNF169 | . | + | 0.181686 | 0.197448 | 1 | hsa_circ_0006705 |
| chr13 | 96636057 | 96651561 | UGGT2 | . | - | 0.181544 | 0.261795 | 1 | hsa_circ_0030632 |
| chr7 | 7826418 | 7841374 | UMAD1 | . | + | 0.176017 | 1.403704 | 0 | hsa_circ_0001676 |
| chr1 | 1747194 | 1770677 | GNB1 | . | - | 0.175834 | 0.186662 | 1 | hsa_circ_0008702 |
| chr2 | 128750760 | 128754065 | SAP130 | . | - | 0.17572 | 0.874603 | 1 | hsa_circ_0056390 |
| chr7 | 158580694 | 158591763 | ESYT2 | . | - | 0.174556 | 0.4188 | 1 | hsa_circ_0001777 |
| chr7 | 138951078 | 138957186 | UBN2 | . | + | 0.174538 | 0.237555 | 1 | hsa_circ_0005594 |
| chr9 | 33953282 | 33996331 | UBAP2 | . | - | 0.173257 | 0.18367 | 1 | hsa_circ_0086720 |
| chr9 | 5954015 | 5988545 | KIAA2026 | . | - | 0.17299 | 0.123703 | 1 | hsa_circ_0138870 |
| chr5 | 70805291 | 70806978 | BDP1 | . | + | 0.171917 | 0.120783 | 1 | hsa_circ_0072897 |
| chr17 | 71231614 | 71233134 | C17orf80 | . | + | 0.168564 | 0.440753 | 1 | hsa_circ_0045537 |
| chr1 | 28362054 | 28384605 | EYA3 | . | - | 0.167732 | 0.092645 | 1 | hsa_circ_0007895 |
| chr2 | 40655612 | 40657441 | SLC8A1 | . | - | 0.167231 | 0.189182 | 0 | hsa_circ_0005232 |
| chr10 | 70404454 | 70406762 | TET1 | . | + | 0.166975 | 2 | 1 | hsa_circ_0093996 |
| chr9 | 37126308 | 37147442 | ZCCHC7 | . | + | 0.165855 | 0.157325 | 0 | hsa_circ_0138745 |
| chr8 | 131164981 | 131193126 | ASAP1 | . | - | 0.163679 | 0.388769 | 1 | hsa_circ_0008934 |
| chr15 | 59179173 | 59179739 | SLTM | . | - | 0.162576 | 0.043467 | 1 | hsa_circ_0000604 |
| chr5 | 38523520 | 38530768 | LIFR | . | - | 0.162546 | 0.453991 | 1 | hsa_circ_0072309 |
| chr9 | 4860124 | 4860901 | RCL1 | . | + | 0.161963 | 0.722902 | 0 | hsa_circ_0007592 |
| chr2 | 190593385 | 190609514 | ANKAR | . | + | 0.158239 | 1.167338 | 1 | hsa_circ_0005644 |
| chr6 | 13579682 | 13584457 | SIRT5 | . | + | 0.156335 | 1.275 | 1 | hsa_circ_0007218 |
| chr5 | 122881110 | 122911657 | CSNK1G3 | . | + | 0.153092 | 0.15686 | 1 | hsa_circ_0073706 |
| chr12 | 19615443 | 19626289 | AEBP2 | . | + | 0.150553 | 0.130873 | 1 | hsa_circ_0006420 |
| chr12 | 26703179 | 26755639 | ITPR2 | . | - | 0.13506 | 0.036917 | 1 | hsa_circ_0098219 |

**Supplementary Table 4.** The 148 circRNAs significantly downregulated in the lesional- relative to non-lesional skin.

| **chromosome** | **start** | **end** | **sense gene** | **antisense gene** | **strand** | **Fold change (RPM)** | ***P*-value** |
| --- | --- | --- | --- | --- | --- | --- | --- |
| chr2 | 99802639 | 99812219 | MRPL30 | . | + | 0.158637 | 0.000166 |
| chr1 | 52981562 | 52992045 | ZCCHC11 | . | - | 0.243481 | 0.000258 |
| chr3 | 171965322 | 171969331 | FNDC3B | . | + | 0.21579 | 0.000312 |
| chrX | 10031484 | 10066619 | WWC3 | . | + | 0.139079 | 0.000353 |
| chr2 | 68717321 | 68772444 | APLF | . | + | 0.09478 | 0.00039 |
| chr8 | 27151596 | 27151827 | TRIM35 | . | - | 0.196087 | 0.000897 |
| chr6 | 57058639 | 57075243 | RAB23 | . | - | 0.213965 | 0.000968 |
| chr15 | 57743697 | 57754090 | CGNL1 | . | + | 0.065762 | 0.001093 |
| chr1 | 155644800 | 155649303 | YY1AP1 | . | - | 0.233373 | 0.001505 |
| chr5 | 137320945 | 137324004 | FAM13B | . | - | 0.172099 | 0.00153 |
| chr8 | 48308935 | 48320523 | SPIDR | . | + | 0.13422 | 0.001989 |
| chr10 | 105767934 | 105778666 | SLK | . | + | 0.238979 | 0.002089 |
| chr2 | 239090705 | 239093928 | ILKAP | . | - | 0.158221 | 0.002244 |
| chrX | 53430497 | 53430825 | SMC1A | . | - | 0.096104 | 0.002504 |
| chr7 | 65705311 | 65751696 | TPST1 | . | + | 0.065237 | 0.002713 |
| chr2 | 40655612 | 40657444 | SLC8A1 | . | - | 0.196529 | 0.00274 |
| chr16 | 18852886 | 18856973 | SMG1 | . | - | 0.278891 | 0.002779 |
| chr17 | 65941524 | 65972074 | BPTF | . | + | 0.267451 | 0.002951 |
| chr21 | 37711076 | 37717005 | MORC3 | . | + | 0.336675 | 0.002976 |
| chr18 | 9182379 | 9221997 | ANKRD12 | . | + | 0.368209 | 0.003201 |
| chr6 | 76344422 | 76388643 | SENP6 | . | + | 0.162774 | 0.003328 |
| chr2 | 168920009 | 168986268 | STK39 | . | - | 0.218139 | 0.00379 |
| chr15 | 41988272 | 41991357 | MGA | . | + | 0.351734 | 0.004125 |
| chr14 | 23375403 | 23380612 | RBM23 | . | - | 0.210776 | 0.004296 |
| chr2 | 113057425 | 113082774 | ZC3H6 | . | + | 0.067063 | 0.004406 |
| chr11 | 107916996 | 107925682 | CUL5 | . | + | 0.252205 | 0.004408 |
| chr9 | 37126308 | 37126939 | ZCCHC7 | . | + | 0.173684 | 0.004743 |
| chr14 | 31602443 | 31602881 | HECTD1 | . | - | 0.164158 | 0.00526 |
| chr22 | 22153300 | 22162135 | MAPK1 | . | - | 0.047042 | 0.005273 |
| chr16 | 53288349 | 53308214 | CHD9 | . | + | 0.02978 | 0.00545 |
| chr7 | 99090662 | 99092254 | ZNF394 | . | - | 0.137221 | 0.005473 |
| chr5 | 63621012 | 63626233 | RNF180 | . | + | 0.110593 | 0.006011 |
| chr2 | 72945231 | 72960247 | EXOC6B | . | - | 0.27868 | 0.006132 |
| chr5 | 132227855 | 132228810 | AFF4 | . | - | 0.28132 | 0.006145 |
| chr10 | 34558584 | 34573173 | PARD3 | . | - | 0.295454 | 0.006753 |
| chr3 | 56626997 | 56628056 | CCDC66 | . | + | 0.279884 | 0.006909 |
| chr13 | 51501542 | 51523641 | RNASEH2B | . | + | 0.288053 | 0.007006 |
| chrX | 139865339 | 139866824 | . | CDR1 | + | 0.38498 | 0.007787 |
| chr2 | 99786012 | 99787892 | MITD1 | . | - | 0.251155 | 0.007985 |
| chr2 | 113057425 | 113089859 | ZC3H6 | . | + | 0.265204 | 0.008096 |
| chr5 | 65349233 | 65350779 | ERBIN | . | + | 0.312715 | 0.008116 |
| chr17 | 57430575 | 57430887 | YPEL2 | . | + | 0.318023 | 0.008185 |
| chr19 | 24014398 | 24016323 | . | . | + | 0.225094 | 0.009631 |
| chr1 | 117944807 | 117963271 | MAN1A2 | . | + | 0.443954 | 0.010449 |
| chr4 | 48371865 | 48396670 | SLAIN2 | . | + | 0.114086 | 0.010778 |
| chr1 | 231930987 | 231954263 | TSNAX-DISC1 | DISC2 | + | 0.098933 | 0.011028 |
| chr1 | 108690900 | 108703915 | SLC25A24 | . | - | 0.077645 | 0.011564 |
| chr2 | 231307651 | 231314970 | SP100 | . | + | 0.227551 | 0.012922 |
| chr4 | 144464661 | 144465125 | SMARCA5 | . | + | 0.402984 | 0.013032 |
| chr8 | 37971709 | 37976881 | ASH2L | . | + | 0.188184 | 0.013276 |
| chr11 | 35222628 | 35229753 | CD44 | . | + | 0.228828 | 0.013363 |
| chr6 | 18236682 | 18258636 | DEK | . | - | 0.43995 | 0.013364 |
| chr10 | 91511102 | 91522592 | KIF20B | . | + | 0.18781 | 0.013409 |
| chr16 | 11114049 | 11154879 | CLEC16A | . | + | 0.196385 | 0.013517 |
| chr9 | 33953282 | 33996331 | UBAP2 | . | - | 0.120549 | 0.013568 |
| chr4 | 48371865 | 48385801 | SLAIN2 | . | + | 0.401495 | 0.014529 |
| chr12 | 46319924 | 46322642 | SCAF11 | . | - | 0.259289 | 0.015524 |
| chr13 | 24164288 | 24200931 | TNFRSF19 | . | + | 0.118221 | 0.015947 |
| chr19 | 47767859 | 47768203 | CCDC9 | . | + | 0.319608 | 0.016108 |
| chr1 | 117944807 | 117957453 | MAN1A2 | . | + | 0.372196 | 0.016328 |
| chr3 | 107429298 | 107435696 | BBX | . | + | 0.19785 | 0.016587 |
| chr8 | 109462051 | 109462721 | EMC2 | . | + | 0.189894 | 0.016803 |
| chr20 | 2944917 | 2945848 | VPS16 | . | + | 0.22606 | 0.017504 |
| chr13 | 26974589 | 26975761 | CDK8 | . | + | 0.112759 | 0.017531 |
| chr7 | 7826418 | 7841374 | UMAD1 | . | + | 0.079041 | 0.017729 |
| chr4 | 37633006 | 37640126 | RELL1 | . | - | 0.337079 | 0.018887 |
| chr1 | 41536266 | 41541123 | SCMH1 | . | - | 0.364579 | 0.018919 |
| chr6 | 158733082 | 158735300 | TULP4 | . | + | 0.334615 | 0.01895 |
| chr4 | 129913321 | 129925031 | SCLT1 | . | - | 0.163804 | 0.019011 |
| chr14 | 39648294 | 39648666 | PNN | . | + | 0.394907 | 0.019137 |
| chr15 | 41961025 | 41962156 | MGA | . | + | 0.267119 | 0.019736 |
| chr18 | 54423813 | 54426184 | WDR7 | . | + | 0.292228 | 0.019968 |
| chr8 | 52773404 | 52773806 | PCMTD1 | . | - | 0.359341 | 0.020024 |
| chr14 | 31416295 | 31425448 | STRN3 | . | - | 0.2215 | 0.020064 |
| chr1 | 33760537 | 33760906 | ZNF362 | . | + | 0.31264 | 0.0201 |
| chr8 | 618597 | 624047 | ERICH1 | . | - | 0.10557 | 0.020737 |
| chr12 | 116668337 | 116675510 | MED13L | . | - | 0.151915 | 0.020759 |
| chr3 | 169863210 | 169867032 | PHC3 | . | - | 0.232745 | 0.020813 |
| chr21 | 30693541 | 30702014 | BACH1 | . | + | 0.267912 | 0.020951 |
| chr13 | 77779390 | 77818086 | MYCBP2 | . | - | 0.168526 | 0.020959 |
| chr7 | 22999874 | 23030758 | FAM126A | . | - | 0.182372 | 0.021152 |
| chr4 | 39739039 | 39776553 | UBE2K | . | + | 0.248604 | 0.021704 |
| chr1 | 243579003 | 243589860 | SDCCAG8 | . | + | 0.216682 | 0.022015 |
| chr20 | 57014000 | 57016139 | VAPB | . | + | 0.161581 | 0.022133 |
| chr11 | 68318588 | 68331900 | PPP6R3 | . | + | 0.130745 | 0.022305 |
| chr1 | 67423741 | 67428843 | MIER1 | . | + | 0.197706 | 0.022483 |
| chr22 | 32874967 | 32875262 | FBXO7 | . | + | 0.172192 | 0.022781 |
| chr3 | 157839891 | 157841780 | RSRC1 | . | + | 0.274643 | 0.023306 |
| chr8 | 141856358 | 141874498 | PTK2 | . | - | 0.291872 | 0.023584 |
| chr9 | 128419929 | 128434922 | MAPKAP1 | . | - | 0.333359 | 0.023621 |
| chr15 | 52161413 | 52194233 | TMOD3 | . | + | 0.234319 | 0.023894 |
| chr13 | 96409897 | 96416207 | DNAJC3 | . | + | 0.421991 | 0.02479 |
| chr6 | 161455290 | 161471011 | MAP3K4 | . | + | 0.363837 | 0.024875 |
| chr15 | 41036244 | 41037457 | RMDN3 | . | - | 0.215814 | 0.025762 |
| chr4 | 148778703 | 148803083 | ARHGAP10 | . | + | 0.234587 | 0.025801 |
| chrX | 84558411 | 84563222 | POF1B | . | - | 0.218827 | 0.026225 |
| chr12 | 123983090 | 123984083 | RILPL1 | . | - | 0.218699 | 0.026724 |
| chr22 | 22160138 | 22162135 | MAPK1 | . | - | 0.350046 | 0.026799 |
| chrX | 84600865 | 84622771 | POF1B | . | - | 0.125441 | 0.0274 |
| chr5 | 82832825 | 82838087 | VCAN | . | + | 0.27867 | 0.027464 |
| chr10 | 112723882 | 112745523 | SHOC2 | . | + | 0.28109 | 0.027503 |
| chr19 | 13039155 | 13039661 | FARSA | . | - | 0.273968 | 0.028596 |
| chr7 | 104714065 | 104717880 | KMT2E | . | + | 0.290637 | 0.030197 |
| chr1 | 174241551 | 174274265 | RABGAP1L | . | + | 0.192876 | 0.030586 |
| chr7 | 155457868 | 155473602 | RBM33 | . | + | 0.333156 | 0.031963 |
| chr20 | 34317233 | 34320057 | RBM39 | . | - | 0.148175 | 0.032057 |
| chr10 | 70404454 | 70406762 | TET1 | . | + | 0.035218 | 0.032208 |
| chr9 | 79996891 | 80022523 | VPS13A | . | + | 0.145651 | 0.032318 |
| chr15 | 64791491 | 64792365 | ZNF609 | . | + | 0.437626 | 0.032589 |
| chr13 | 41515056 | 41518061 | ELF1 | . | - | 0.372304 | 0.032631 |
| chr8 | 142264087 | 142264728 | SLC45A4 | LOC105375787 | - | 0.421744 | 0.03321 |
| chr14 | 31398406 | 31425448 | STRN3 | . | - | 0.031687 | 0.033596 |
| chr17 | 45479497 | 45492285 | EFCAB13 | . | + | 0.18208 | 0.033753 |
| chr12 | 109046047 | 109048186 | CORO1C | . | - | 0.328334 | 0.034236 |
| chr8 | 42761315 | 42798588 | HOOK3 | . | + | 0.263237 | 0.034714 |
| chr15 | 44624185 | 44630515 | CASC4 | . | + | 0.124337 | 0.035607 |
| chr14 | 67736417 | 67770316 | MPP5 | . | + | 0.094323 | 0.03567 |
| chr19 | 33604672 | 33605325 | GPATCH1 | . | + | 0.441829 | 0.036087 |
| chr14 | 31404368 | 31425448 | STRN3 | . | - | 0.273492 | 0.036103 |
| chr2 | 24357988 | 24369956 | FAM228B | . | + | 0.126438 | 0.036447 |
| chr8 | 18656804 | 18662408 | PSD3 | . | - | 0.300001 | 0.03691 |
| chr21 | 17205666 | 17214859 | USP25 | . | + | 0.293932 | 0.0374 |
| chr11 | 35222628 | 35226187 | CD44 | . | + | 0.434217 | 0.037642 |
| chr9 | 33986757 | 34017187 | UBAP2 | . | - | 0.250713 | 0.038564 |
| chr4 | 129857809 | 129891623 | SCLT1 | . | - | 0.115849 | 0.03982 |
| chr2 | 61343113 | 61345251 | KIAA1841 | . | + | 0.299925 | 0.039985 |
| chr15 | 63988322 | 64008672 | HERC1 | . | - | 0.332667 | 0.040135 |
| chr7 | 155465560 | 155473602 | RBM33 | . | + | 0.430998 | 0.040153 |
| chr1 | 117944807 | 117948267 | MAN1A2 | . | + | 0.408137 | 0.040647 |
| chr9 | 138773478 | 138774924 | CAMSAP1 | . | - | 0.468012 | 0.040651 |
| chr7 | 30590251 | 30614497 | LOC401320 | . | - | 0.330021 | 0.041221 |
| chr1 | 8601272 | 8617582 | RERE | . | - | 0.324521 | 0.041543 |
| chr3 | 3178943 | 3186394 | TRNT1 | . | + | 0.218829 | 0.041728 |
| chr1 | 117944807 | 117984947 | MAN1A2 | . | + | 0.394201 | 0.042464 |
| chr11 | 35222628 | 35231601 | CD44 | . | + | 0.3493 | 0.043727 |
| chr5 | 70805291 | 70806978 | BDP1 | . | + | 0.254608 | 0.043942 |
| chr2 | 168869143 | 168873605 | STK39 | . | - | 0.33644 | 0.043969 |
| chr3 | 138289159 | 138291774 | CEP70 | . | - | 0.355982 | 0.044328 |
| chr5 | 80071512 | 80088663 | MSH3 | . | + | 0.368446 | 0.045042 |
| chr9 | 19286766 | 19305525 | DENND4C | . | + | 0.113402 | 0.045132 |
| chr14 | 102661274 | 102676199 | WDR20 | . | + | 0.146936 | 0.045536 |
| chr8 | 18622958 | 18662408 | PSD3 | . | - | 0.228183 | 0.045718 |
| chr1 | 67356836 | 67371058 | WDR78 | . | - | 0.200091 | 0.047252 |
| chr2 | 190593385 | 190609514 | ANKAR | . | + | 0.186224 | 0.047417 |
| chr7 | 158580694 | 158591763 | ESYT2 | . | - | 0.221588 | 0.047924 |
| chr2 | 160194077 | 160252345 | BAZ2B | . | - | 0.175174 | 0.048943 |
| chr3 | 169854206 | 169867032 | PHC3 | . | - | 0.394944 | 0.04908 |
| chr5 | 167915606 | 167921655 | RARS | . | + | 0.223233 | 0.049101 |

**Supplementary Table 5.** The 37 significantly differentially expressed miRNAs among the 137 high abundance miRs.

| **miRNA** | **P value** | **Mean Lesional** | **Mean Non-lesional** | **Fold change** |
| --- | --- | --- | --- | --- |
| hsa-miR-4454+hsa-miR-7975 | 0.003602 | 4053.99 | 385.405 | 10.51877895 |
| hsa-miR-205-5p | 0.002738 | 3895.7 | 1729.63 | 2.252331423 |
| hsa-miR-203a-3p | 0.003996 | 387.687 | 41.58 | 9.323881674 |
| hsa-let-7g-5p | 0.012657 | 985.67 | 663.535 | 1.485483057 |
| hsa-let-7i-5p | 0.011747 | 732.137 | 410.588 | 1.783142712 |
| hsa-miR-223-3p | 0.046486 | 210.042 | 1.51 | 139.1006623 |
| hsa-miR-200c-3p | 0.001915 | 334.265 | 140.808 | 2.373906312 |
| hsa-miR-126-3p | 0.00533 | 226.407 | 55.5567 | 4.07524205 |
| hsa-miR-191-5p | 0.022546 | 296.523 | 144.072 | 2.058158421 |
| hsa-miR-146a-5p | 0.000277 | 133.56 | 1.51 | 88.45033113 |
| hsa-miR-27b-3p | 0.000062 | 128.808 | 15.1467 | 8.504030581 |
| hsa-miR-16-5p | 0.000436 | 92.7967 | 4.92333 | 18.84836076 |
| hsa-let-7f-5p | 0.008361 | 141.24 | 70.26 | 2.010247652 |
| hsa-miR-200b-3p | 0.026893 | 94.995 | 24.3183 | 3.906317465 |
| hsa-miR-25-3p | 0.024448 | 82.1167 | 17.1083 | 4.799816463 |
| hsa-miR-93-5p | 0.001073 | 71.8783 | 7.875 | 9.127403175 |
| hsa-miR-222-3p | 0.047431 | 70.6133 | 24.4067 | 2.893193263 |
| hsa-miR-221-3p | 0.001767 | 44.9483 | 1.51 | 29.76708609 |
| hsa-miR-199a-5p | 0.02715 | 80.6133 | 45.1383 | 1.785917946 |
| hsa-miR-365a-3p+hsa-miR-365b-3p | 0.005039 | 47.9933 | 15.325 | 3.131699837 |
| hsa-miR-21-5p | 0.031252 | 30.755 | 1.51 | 20.36754967 |
| hsa-miR-155-5p | 0.000795 | 20.7517 | 1.51 | 13.74284768 |
| hsa-miR-99b-5p | 0.001419 | 19.445 | 1.51 | 12.87748344 |
| hsa-miR-20a-5p+hsa-miR-20b-5p | 0.028794 | 18.7917 | 1.51 | 12.44483444 |
| hsa-miR-24-3p | 0.031005 | 16.035 | 2.72 | 5.895220588 |
| hsa-miR-148a-3p | 0.029742 | 14.4717 | 1.51 | 9.583907285 |
| hsa-miR-342-3p | 0.049107 | 19.39 | 6.76833 | 2.864813034 |
| hsa-miR-15a-5p | 0.037537 | 13.765 | 1.51 | 9.11589404 |
| hsa-miR-26b-5p | 0.043996 | 13.3733 | 1.51 | 8.856490066 |
| hsa-miR-148b-3p | 0.01364 | 9.75667 | 1.80167 | 5.415347983 |
| hsa-miR-127-3p | 0.036368 | 10.6017 | 2.745 | 3.862185792 |
| hsa-miR-574-5p | 0.017697 | 15.5767 | 54.65 | 0.285026532 |
| hsa-miR-125a-5p | 0.043669 | 131.488 | 173.888 | 0.756164888 |
| hsa-miR-612 | 0.000727 | 27.7233 | 80.6217 | 0.343868958 |
| hsa-miR-302d-3p | 0.031162 | 135.05 | 353.245 | 0.382312559 |
| hsa-let-7c-5p | 0.000364 | 887.303 | 2080.34 | 0.426518261 |
| hsa-let-7b-5p | 0.000356 | 5846.61 | 13321.4 | 0.438888555 |
